# The *Pseudomonas aeruginosa* accessory genome elements influence virulence towards *Caenorhabditis elegans*

**DOI:** 10.1101/621433

**Authors:** Alejandro Vasquez-Rifo, Isana Veksler-Lublinsky, Zhenyu Cheng, Frederick M. Ausubel, Victor Ambros

**Affiliations:** Program in Molecular Medicine, University of Massachusetts Medical School, Worcester, Massachusetts, 01605, USA; Department of Software and Information Systems Engineering, Ben-Gurion University of the Negev, Beer-Sheva, Israel; Department of Microbiology & Immunology, Dalhousie University, Halifax, Nova Scotia, Canada; Department of Molecular Biology, Massachusetts General Hospital, Boston, MA 02114 and Department of Genetics, Harvard Medical School, Boston, MA 02115

**Keywords:** *C. elegans*, *P. aeruginosa*, accessory genome, virulence, CRISPR-Cas

## Abstract

Multicellular animals and bacteria frequently engage in predator-prey and host-pathogen interactions, such as the well-studied relationship between *Pseudomonas aeruginosa* and the nematode *Caenorhabditis elegans*. This study investigates the genomic and genetic basis of bacterial-driven variability in *P. aeruginosa* virulence towards *C. elegans*. Natural isolates of *P. aeruginosa* that exhibit diverse genomes display a broad range of virulence towards *C. elegans*. Using gene association and genetic analysis, we identified accessory genome elements that correlate with virulence, including both known and novel virulence determinants. Among the novel genes, we found a viral-like mobile element, the *teg* block, that impairs virulence and whose acquisition is restricted by CRISPR-Cas systems. Further genetic and genomic evidence suggests that spacer-targeted elements preferentially associate with lower virulence and suggest a positive, albeit indirect, role for host CRISPR-Cas systems in the restriction of accessory genome elements that may be detrimental to virulence.

## INTRODUCTION

Interactions between environmental bacteria and small invertebrate animals, such as free-living nematodes, are ecologically significant in many terrestrial ecosystems (Ferris, 2010). These interactions comprise many types of ecological relationships that range from reciprocal harm to mutualism. Frequently, animal-bacterial interactions are ‘predator-prey’ relationships, where for example nematodes feed on bacteria. Such predation can in turn drive the evolution of bacterial anti-predator mechanisms, such as the production of noxious toxins, and/or full pathogenic potential where the bacterium can kill and feed on the predator ((Weitere et al., 2005); reviewed in (Jousset, 2012)). One such bacterial species is *Pseudomonas aeruginosa* (*P. aeruginosa*), that is preyed upon by invertebrates and is also a facultative pathogen of a broad range of hosts including plants, amoeboid protists, insects, mammals, and nematodes (Mahajan-Miklos et al., 1999; Pukatzki et al., 2002; Rahme et al., 1995, 1997).

The relationship between a facultatively pathogenic bacterium and a predator, such as a free-living nematode, can be bidirectional, with the pathogen either serving as a food source for the predator, or itself thriving on the infected predator. For example, the nematode *Caenorhabditis elegans* (*C. elegans*) (Weitere et al., 2005) can grow from larval stages to the adult by feeding on the pathogenic bacterium *P. aeruginosa*. Interestingly, although *C. elegans* larval development can proceed successfully on *P. aeruginosa*, adults can suffer dramatically reduced lifetimes, depending on the *P. aeruginosa* strain (for example, median adult survival of ∼2 days on strain PA14 compared to ∼14 days on *Escherichia coli* strain OP50 that is used as standard diet). This mutually-antagonistic relationship between *C. elegans* and *P. aeruginosa* is a well-studied model for ecologically coexisting predators of *P. aeruginosa* that are also natural hosts for infection (Tan et al., 1999).

It is plausible that *C. elegans* and *P. aeruginosa* interact in natural niches, as the bacterium is known to inhabit many environments including soils (Deredjian et al., 2014; Kaszab et al., 2011; Rutherford et al., 2018) and the nematode is often an inhabitant of soil and rotting plant matter (Schulenburg and Félix, 2017). These interactions could be transitory in the wild, due to worm avoidance of *P. aeruginosa* or death of the worms, and thus difficult to catalog, but have been sustained by a report of natural coexistence of the two species (Grewal PS 1991, reviewed in (Schulenburg and Félix, 2017)).

In the present work, we addressed the sources and genomic correlates of variability in the virulence of distinct *P. aeruginosa* strains towards *C. elegans*. A previous study of 20 *P. aeruginosa* natural isolates revealed strain-driven variation in *P. aeruginosa* virulence, highlighting virulence as a complex trait, likely the result of multiple components acting in a combinatorial manner (Lee et al., 2006). Extending this previous work, we conducted an in-depth genome-wide comparative survey of a set of 52 *P. aeruginosa* strains. We used comparative genomic approaches to identify correlations between *P. aeruginosa* virulence and the presence/absence of specific accessory genome elements, including bacterial immune defense systems.

Our analysis revealed gene sets in the accessory genome of *P. aeruginosa* (*i.e*. the set of genes present in some, but not all, of the strains in the species) that correlate either with high or low virulence. Our approach identified known virulence factors, as well as novel factors that can directly modulate bacterial virulence, either positively or negatively. We also identified genes that may indirectly affect virulence. For example, our study revealed a positive role in virulence for certain bacterial immune defense systems which filter horizontal gene transfer (HGT), and hence can impact the composition of the accessory genome. In particular, we found that *P. aeruginosa* strains with active CRISPR-Cas systems have statistically higher levels of virulence towards *C. elegans* and that spacer-targeted genes are among the genes associated with lower virulence. These correlative findings, together with our genetic confirmation of virulence-inhibitory activity of certain accessory genome elements, support an indirect role for CRISPR-Cas systems in contributing to the maintenance and evolution of high virulence against nematodes.

## RESULTS

### A large *P. aeruginosa* accessory genome underlies substantial strain diversity in gene content

To assess the extent of variation in genetic makeup among a diverse panel of environmental and clinical *P. aeruginosa* strains, we analyzed *in silico* the genomes of 1488 *P. aeruginosa* strains. The protein-coding genes of the strains were assigned to clusters of homologous genes using the CD-HIT program (Fu et al., 2012) with a threshold of 70% amino acid similarity. The clustering procedure resulted in the identification of 28,793 distinct gene clusters (i.e. groups of homologous genes). We then examined the distribution and frequency of these 28,793 genes across the 1,488 *P. aeruginosa* strains. 5,170 genes were present in more than 90% of the isolates and were accordingly defined as constituting the *P. aeruginosa* core genome (Figure 1A). The remaining 23,623 genes constitute the accessory genome of these 1,488 *P. aeruginosa* strains. The frequency distribution of the genes is bimodal, with prominent maxima corresponding to the core genome and the set of genes that occur only once in these strains (referred to as ‘singletons’, Figure 1B). The ratio between the pangenome and the core genome (5.6) agrees with a previously reported ratio: 5.3 (van Belkum et al., 2015), confirming that *P. aeruginosa* harbors a large amount of strain-specific variation in protein-coding genes.

**Figure 1.**
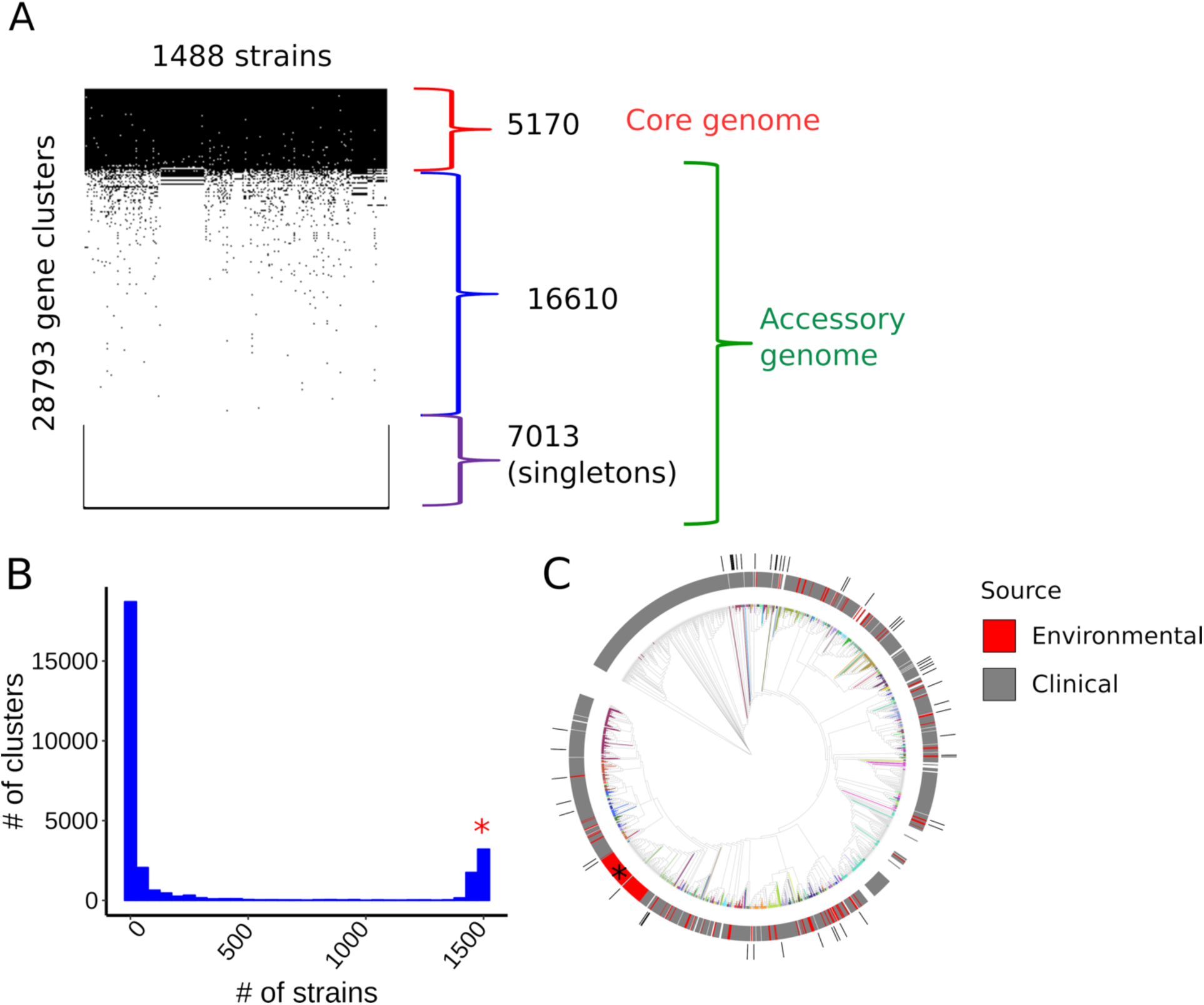
Pangenomic and phylogenetic features of *Pseudomonas aeruginosa*. **A)** Presence/Absence matrix of *P. aeruginosa* genes across the strains. Core and accessory genomes are marked. **B**) The frequency distribution of the genes among the 1488 strains. The right end of the distribution, marked with red *, corresponds to the core genome, while the left end of the distribution, corresponds to singletons and rare accessory genes. **C)** Phylogenetic tree of *P. aeruginosa* strains. Phylogenetically related MLST groups are shown in different colors. Isolation source is shown on top of the tree. An apparent clade enriched for environmental strains (indicated by *) is artificially enlarged by the repeated presence of a set of almost identical genomes in the set used to build the phylogeny. The phylogenetic locations of the 52 isolates experimentally tested in this study are indicated in the outer circle (black bars).

To model the phylogenetic relationships between the *P. aeruginosa* isolates, we aligned the core genomes and used the alignments to build a phylogenetic tree (Figure 1C). The isolation source of the strains, when available, was categorized as clinical or environmental and this designation was mapped to the tree (Figure 1C). Environmental strains distribute widely across the tree and do not associate with any clade in particular. The result is consistent with other studies that showed that both clinical and environmental isolates of *P. aeruginosa* can originate from the same clade (Kidd et al., 2012; Pirnay et al., 2005, 2009; Selezska et al., 2012).

In order to experimentally study the effect of bacterial genetic variation on the interaction between *P. aeruginosa* and *C. elegans*, we assembled a collection of 52 representative *P. aeruginosa* strains (Supplemental Table 1) included in the *in silico* collection of 1,488. The collection consists of bacterial isolates derived from clinical (85%, mostly from primary infections) and environmental (15%) settings. The 52 strains distributed widely across *P. aeruginosa* phylogeny, with no particular bias towards any specific clade (Figure 1C). The 52-strain cohort have a pangenome of 11,731 genes and an accessory genome of 6,537 genes.

### Virulence towards the nematode *C. elegans* strongly varies among *P. aeruginosa* strains

To assess phenotypic variation in interactions of *P. aeruginosa* with *C. elegans*, we measured the virulence towards *C. elegans* wildtype worms for the collection of 52 *P. aeruginosa* strains. Young adult *C. elegans* hermaphrodites were exposed to a full lawn of each *P. aeruginosa* strain using so-called slow kill (SK) media (Tan et al., 1999). These assay conditions minimize the effects of worm behavior on survival (Martin et al., 2017; Reddy et al., 2009) and promote bacterial colonization of the worm gut (Tan et al., 1999). Adult lifetime was scored using a semi-automated method (Stroustrup et al., 2013) to obtain survival curves for worms exposed to each bacterial strain (Figure 2A). Bacterial strain virulence towards *C. elegans* was measured as the median survival time of worms exposed to each bacterial strain (Figure 2B). Virulence varied continuously over a five-fold range, spanning from 1.5 days to over 10 days (Figure 2B). Indeed, the median worm survival on *P. aeruginosa* for strain z7, which exhibited the lowest virulence towards *C. elegans*, was greater than that of worms exposed to *E. coli* HB101, a strain commonly used in the laboratory to maintain worm stocks (Figure 2B). In addition, under SK conditions, the number of viable progeny produced by hermaphrodites exposed to strain z7 was indistinguishable from that of animals exposed to *E. coli* HB101 (Supplemental Figure 1A). Altogether, these results show that for our experimental set of 52 *P. aeruginosa* strains, virulence varies continuously over a wide range, from highly virulent strains, which kill *C. elegans* adults within 2 days, to essentially completely avirulent strains that do not detectably impair worm lifespan or reproduction in comparison to their normal laboratory food.

**Figure 2.**
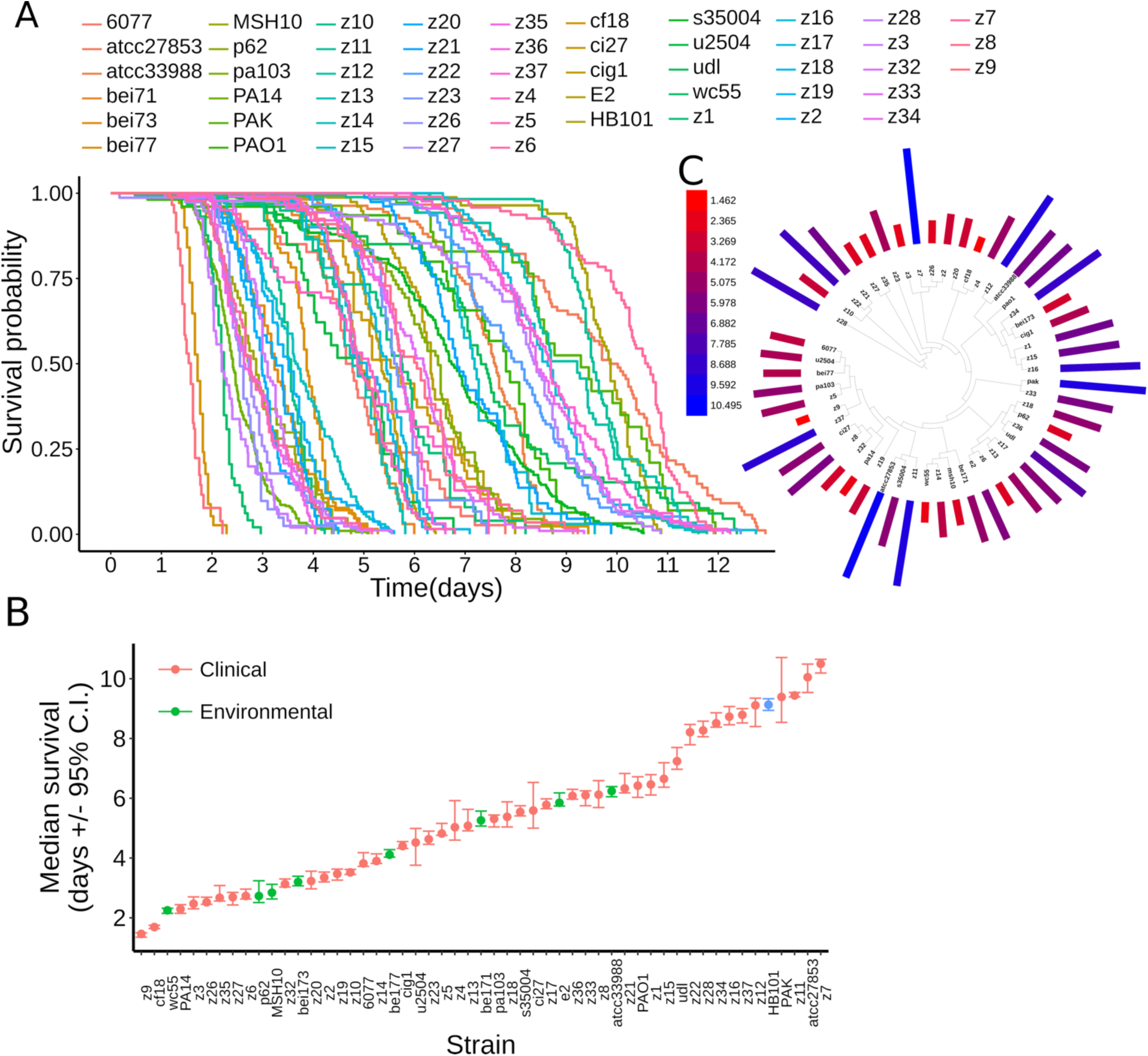
*P. aeruginosa* virulence towards adult *C. elegans* worms. **A**) Survival curves of adult *C. elegans* worms exposed to the studied collection of 52 *P. aeruginosa* strains. **B**) Median survival of adult *C. elegans* worms exposed to the studied collection of *P. aeruginosa* strains (left panel, confidence interval, C.I.). The source of the strains is categorized as clinical (colored red) or environmental (colored green). The *E. coli* strain HB101 is included as comparative control and is colored blue. **C**) Phylogenetic distribution of virulence. The virulence of each isolate (i.e. median worm survival in days) is indicated by a bar with length proportional to its value and colored in a gradient, as indicated by the heatmap legend (virulence values in days).

To evaluate the potential contribution of strain isolation source to virulence against *C. elegans*, we compared the set of clinical isolates to the environmental isolates. Strains from clinical settings displayed lower mean virulence when compared to strains isolated from non-clinical, environmental settings (Welch *t*-test, p-value = 0.047, Supplemental Figure 1B). This result suggests that clinical strains isolated from infected humans do not constitute a biased sampling of strains that are relatively more pathogenic to worms than environmental isolates. Rather, it is possible that some clinical strains could harbor variations and adaptations that disfavor virulence towards worms.

Next, we evaluated the distribution of virulence along the *P. aeruginosa* phylogeny. Mapping of virulence onto the phylogenetic tree of the studied isolates showed no phenotypic clustering of virulence towards any particular clade (Fig. 2C). Thus, evolutionarily fluctuations in virulence among isolates, occur without any particular affiliation to select phylogenetic clades.

Defects in bacterial growth rates can impair virulence towards *C. elegans*, and such impairments can be detected *in vitro* (e.g. Feinbaum et al., 2012). Thus, we assessed whether strain-specific virulence against *C. elegans* could primarily reflect the relative growth rate capacity of each strain, as determined by growth rate in LB media at 25°C (the temperature of the virulence assays). We found that growth rate in LB medium showed no statistically significant correlation with virulence (Supplemental Figure 2, Pearson’s correlation, ρ = −0.3, p-value = 0.08).

### *P. aeruginosa* virulence correlates with the presence of particular accessory genome elements

We employed gene association analysis to test whether virulence of *P. aeruginosa* strains towards *C. elegans* could be associated with the presence or absence of specific bacterial genes. In this analysis, virulence is defined as a quantitative trait for each strain, corresponding to the mean lifespan of adult *C. elegans* hermaphrodites when fed each of the strains. The association between genes and virulence was measured using the Mann-Whitney (MW) and linear regression (LR) tests, followed by a gene permutation approach, to assess the reliability of the p-value. Furthermore, genes with significant associations, as determined by the MW and LR tests, were evaluated with two additional metrics that consider phylogeny to resolve confounding effects due to population structure, namely, the ‘simultaneous’ and ‘subsequent’ scores described by Collins and Didelot (2018) (Supplemental Table 2). Gene associations were assessed for the set of 11,731 protein-coding pangenomic genes of the 52 experimental strains, and for a set of 83 previously-identified non-coding RNA genes (excluding rRNAs and tRNAs) of *P. aeruginosa.*

The small non-coding RNAs of bacteria fulfill diverse gene regulatory roles and can modulate pathways required for virulence (Kay et al., 2006; Zhang et al., 2017). Interestingly, we noted that most of the non-coding RNA genes we examined are core genome elements (78%, 65/83 genes). We found no statistically significant association between the non-coding RNAs of *P. aeruginosa* and virulence (Supplemental Figure 3A, all p-value > 0.05 for the MW and LR tests).

**Figure 3.**
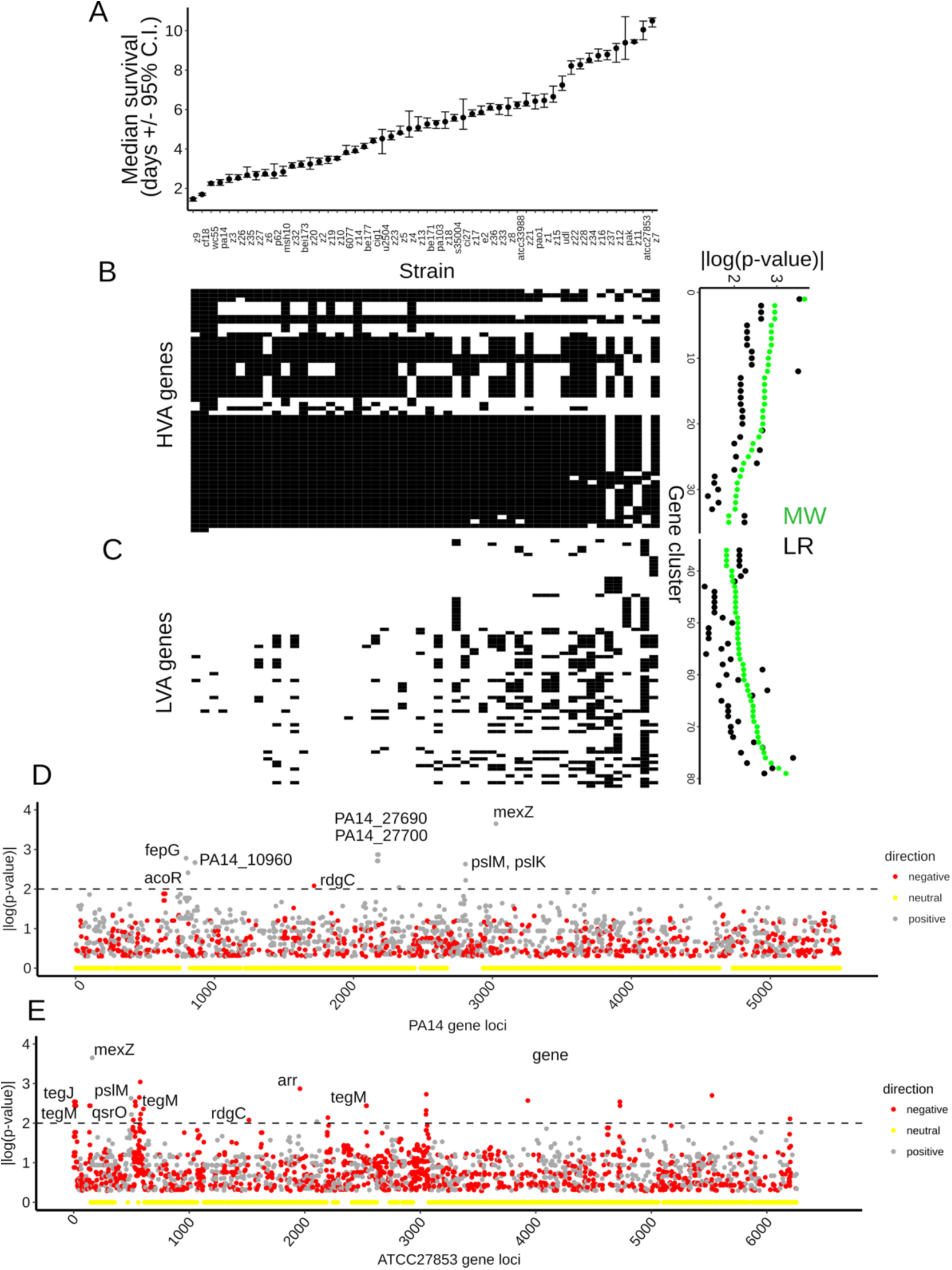
Association between protein-coding genes of *P. aeruginosa* and bacterial virulence. **A**) Median survival of adult *C. elegans* worms exposed to a collection of 52 *P. aeruginosa* strains. The strains are ordered from high to low virulence (left to right) and aligned with the matrixes below. (**B**) Left panels: gene presence/absence matrix for HVA genes (top) and LVA genes (bottom). Gene presence is indicated with black squares and absence with white squares. Genes (rows) are aligned with the corresponding p-values. Right panels: Association statistics (p-value of MW and LR tests) for the HVA and LVA genes, shown as | log_10_(pval)|. (**C-D**) Associated genes present in the strain PA14 (C) or ATCC27853 (D). Gene loci are plotted against the association statistic (p-value of MW test), shown as | log_10_(pval)|. Loci are colored according to the directionality of the gene-virulence association (grey: positively associated; red: negatively associated; yellow: p-value equals zero). Horizontal dashed lines demarcate a significance threshold (p < 0.01).

Among the 6,537 protein-coding accessory genes present in the 52-strain experimental panel, we identified 79 genes significantly associated with virulence, either positively, or negatively (Figure 3, p-value < 0.01 for the MW or LR tests). For 35 of these 79 virulence-associated genes (44%), their presence defined a set of strains with higher virulence compared to the strain set where the same genes were absent (Figure 3B). We refer to them as high virulence-associated genes (or ‘HVA genes’ for short). For the other 44 genes (56%) their presence corresponded to strains with lower virulence (Figure 3B). We refer to these as low virulence-associated genes (or ‘LVA genes’ for short). Each strain harbors a different subset of the 79 associated genes. For example, strain PA14, a highly virulent strain, has 19 HVA genes and 1 LVA gene (Figure 3C). On the other side of the spectrum, strain ATCC27853, a poorly virulent isolate, has 5 HVA genes and 41 LVA genes (Figure 3D). A description of the 79 genes associated with higher or lower virulence is presented in Supplemental Table 2. All the LVA genes (44/44 or 100%) were supported by either the simultaneous or subsequent scores (p-value < 0.05). Similarly, 30/35 of the HVA genes (86%) were supported by either simultaneous or subsequent scores (p-value < 0.05, Supplemental Table 1). Altogether, these phylogenetically-aware scores suggest that population structure does not confound interpretation of the gene associations observed. This result is also congruent with the absence of phenotypic clustering of virulence in the phylogenetic tree (Figure 2D).

The 79 virulence-associated genes encompass a variety of functions, although for many of the associated genes, a functional annotation is not available (43% of HVA genes and 64% of the LVA genes are annotated as ‘hypothetical proteins’). Associated genes could be categorized as follows: 1) Genes with known regulatory roles: Such roles can be ascribed to strain PA14 genes PA14_27700 (HVA gene #13286) and PA14_27690 (HVA gene #15454), which encode a cAMP-dependent protein kinase and RNA polymerase sigma factor, respectively. A second example is the *qsrO* gene (LVA gene #17701), which negatively regulates a highly conserved quorum sensing pathway (Köhler et al., 2014). 2) Genes that encode proteins associated with structural roles: The *pslM* (HVA gene #2628) and *pslK* (HVA gene #2479) genes belong to the psl polysaccharide biosynthetic pathway, a polymer that contributes to biofilm formation (Franklin et al., 2011). Other examples are the HVA genes #6371, #8276 and #8113, which encode homologs of *wbpZ, wbpL and wzz*, respectively. These homologs encode enzymes required for LPS O-antigen synthesis (Rocchetta et al., 1999), a structural component of the bacterial outer membrane. 3) Mobile genetic elements: Several of the genes associated with low virulence are annotated as integrase (genes #6157, #4439, #10878, #8459)”, or phage-related (genes #8274, #5222), suggests that these genes are likely to encode components of mobile genetic elements. Further support for the mobility of these elements comes from their targeting by CRISPR spacers (see below).

Among the genes that we found to be associated with high virulence across the 52-strain panel, two HVA genes, PA14_27700 and PA14_27690, have been previously characterized as virulence genes. Previous genetic analysis showed that loss of function mutations in either PA14_27700 (HVA gene #13286) or PA14_27690 (HVA gene #14622) compromised the virulence of strain PA14 against *C. elegans* (Feinbaum et al., 2012) under the SK assay conditions, the same condition used in the present study. Our examination of the published literature identified a total of 60 previously-described *P. aeruginosa* virulence genes (Supplemental Table 3), that were identified by genetic analysis of virulence against *C. elegans* for two commonly studied *P. aeruginosa* strains, PA14 and PAO1 (Figure 4A-B), both of which are included in our experimental test panel. Upon analysis of these 60 genes, we found that two of the HVA genes associated with virulence in our 52-strain panel (Supplemental Table 2), *pslM* (HVA gene #2628) and *pslK* (HVA gene #2479), were not previously identified as virulence genes in PA14 or PAO1, but are contained in the same *psl* operon as the previously known virulence gene *pslH* (gene #6064), which was shown to be required for full virulence in the PAO1 strain (van Tilburg Bernardes et al., 2017).

**Figure 4.**
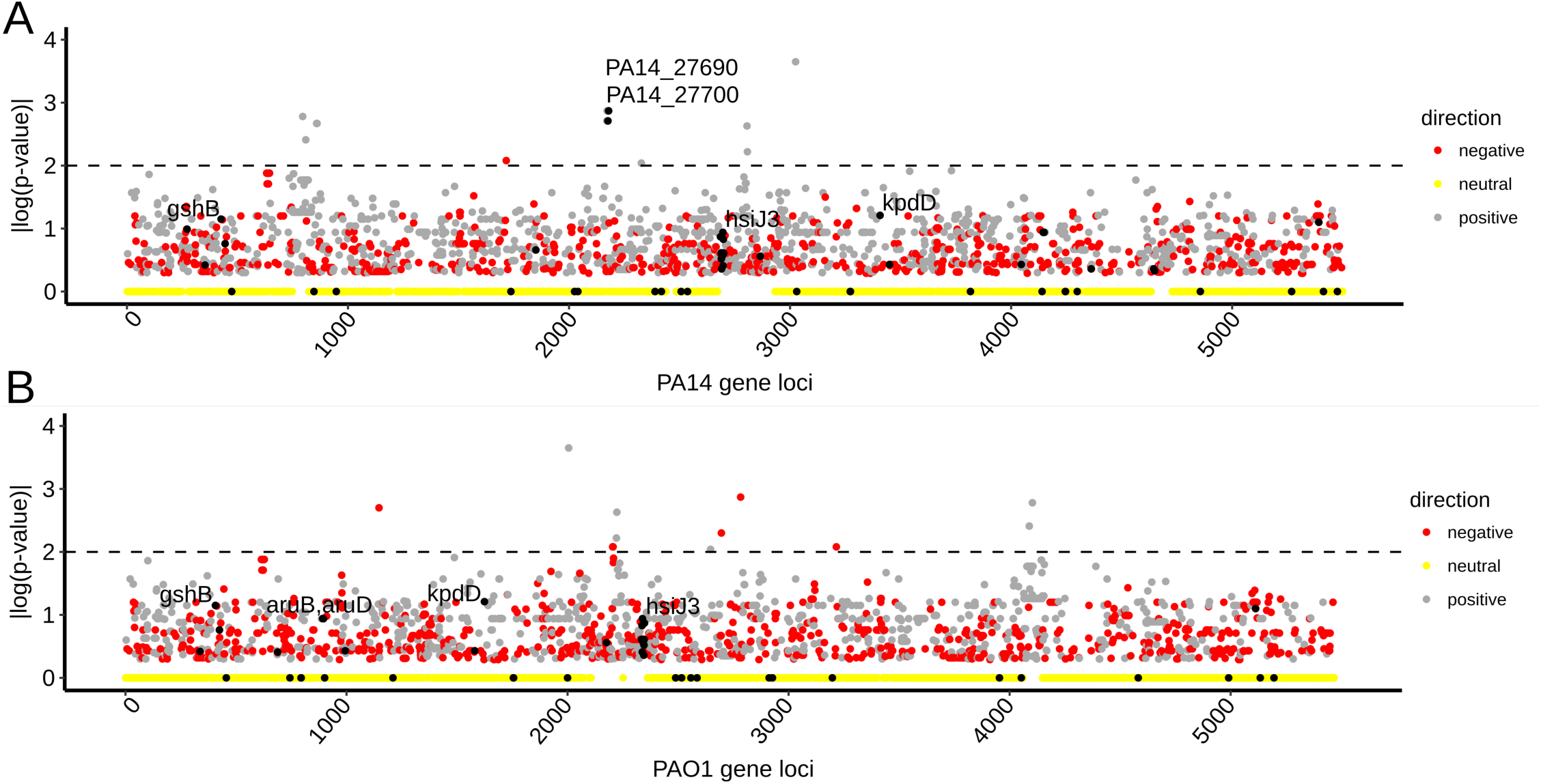
Distribution and features of previously-identified virulence genes. **A-B**) Gene association for PA14 (**A**) and PAO1 (**B**) protein-coding gene loci. Gene loci are plotted against the association statistic (p-value of MW test), shown as | log10(p-value)|. Previously known virulence genes are indicated with black dots and the top 5 most associated genes labelled. The top known genes associated with virulence are PA14_27690 and PA14_27700. Loci are also colored according to the directionality of the gene-virulence association (grey: positively associated; red: negatively associated; yellow: p-value equals zero). Horizontal dashed lines demarcate a significance threshold (p < 0.01).

Other than PA14_27700, PA14_27690 and the *psl* operon genes (*pslM, pslK*), no other genes from the set of 60 previously-described virulence factors showed association with virulence in this study (Figure 4; Supplemental Figure 3B). Notably, 51 of the 60 known virulence genes (85%) belong to the core genome of our panel of 52 experimental strains, explaining the null association observed. The remaining previously-known virulence genes that did not emerge as HVA genes in our 52-strain panel may not have a strong enough impact on virulence across our 52 stains for a variety of potential reasons, including strain-specific epistasis from other accessory genome elements.

### Genetic tests identify *P. aeruginosa* accessory genome elements that contribute to decreased or increased virulence towards *C. elegans*

The statistical association of particular protein-coding genes with either high virulence (in the case of HVA genes) or low virulence (in the case of LVA genes) across the set of 52 experimental strains tested here could in principle reflect the presence or absence of single genes that are individually necessary and/or sufficient to impact virulence. In such cases, loss-of-function or gain-of-function genetic manipulations of the relevant strains would be expected to measurably impact virulence. However, single gene causality may in some cases be masked by strain-specific epistatic interactions, for example with other accessory genes. It would not be unexpected if some of the HVA and LVA genes that we identified were to function in combination, such that the contribution of each individual gene would not be easily evident from single gene knock out or overexpression tests. It is also possible that a gene with no direct function in virulence could nevertheless show association with virulence because of a physiological or ecological linkage between the function of that gene and the function and/or acquisition of *bona fide* virulence factors.

The above expected caveats notwithstanding, we used loss-of-function and gain-of-function approaches to test whether individual HVA genes are necessary and/or sufficient to support high virulence, and conversely, whether LVA genes are necessary and/or sufficient to impose reduced virulence. For most of these genetic tests we selected strain z8, which exhibits an intermediate level or virulence, contains members of both the HVA and LVA gene sets, and is amenable to genome-editing through use of its endogenous CRISPR-Cas system.

The set of HVA genes included previously validated virulence genes (e.g. PA14_27700, PA14_27690), which we did not re-test here. Instead, we evaluated the potential role in virulence for *mexZ* (gene #14466), which had not been previously tested genetically. We constructed an in-frame deletion of *mexZ* in strain z8 (*ΔmexZ*), but no difference in virulence was found for *ΔmexZ* when compared to the wildtype z8 strain (Supplemental Figure 4). The absence of a direct effect on virulence of strain z8 suggests that the association of *mexZ* with virulence amongst the panel of 52 strains could be secondary to additional underlying factors. *mexZ* is frequently mutated in clinical isolates, as a part of the bacterial adaptations to acquire antibiotic resistance (Aires et al., 1999; Westbrock-Wadman et al., 1999).

We next selected genes associated with low virulence to test their effects by using loss of function and gain of function approaches. We assigned gene names to the genes selected for study that were not previously named (Figure 5A and Supplemental Table 4). The selected genes belong to three genomic loci: the *ghlO* gene (LVA gene# 25296) is associated with virulence as a single gene (*i.e*. no additional neighboring genes are associated with virulence); the *qsrO* gene (LVA gene# 17701, (Köhler et al., 2014)) belongs to a four gene operon (referred to as ‘*qsr’* operon); the *tegG* to *tegN* genes (LVA genes # 5222, 5330, 10513, 15466, 21386, 21557, 26140), constitute a block of contiguous genes in bacterial chromosomes (referred to as the ‘*teg* block’, described below).

**Figure 5.**
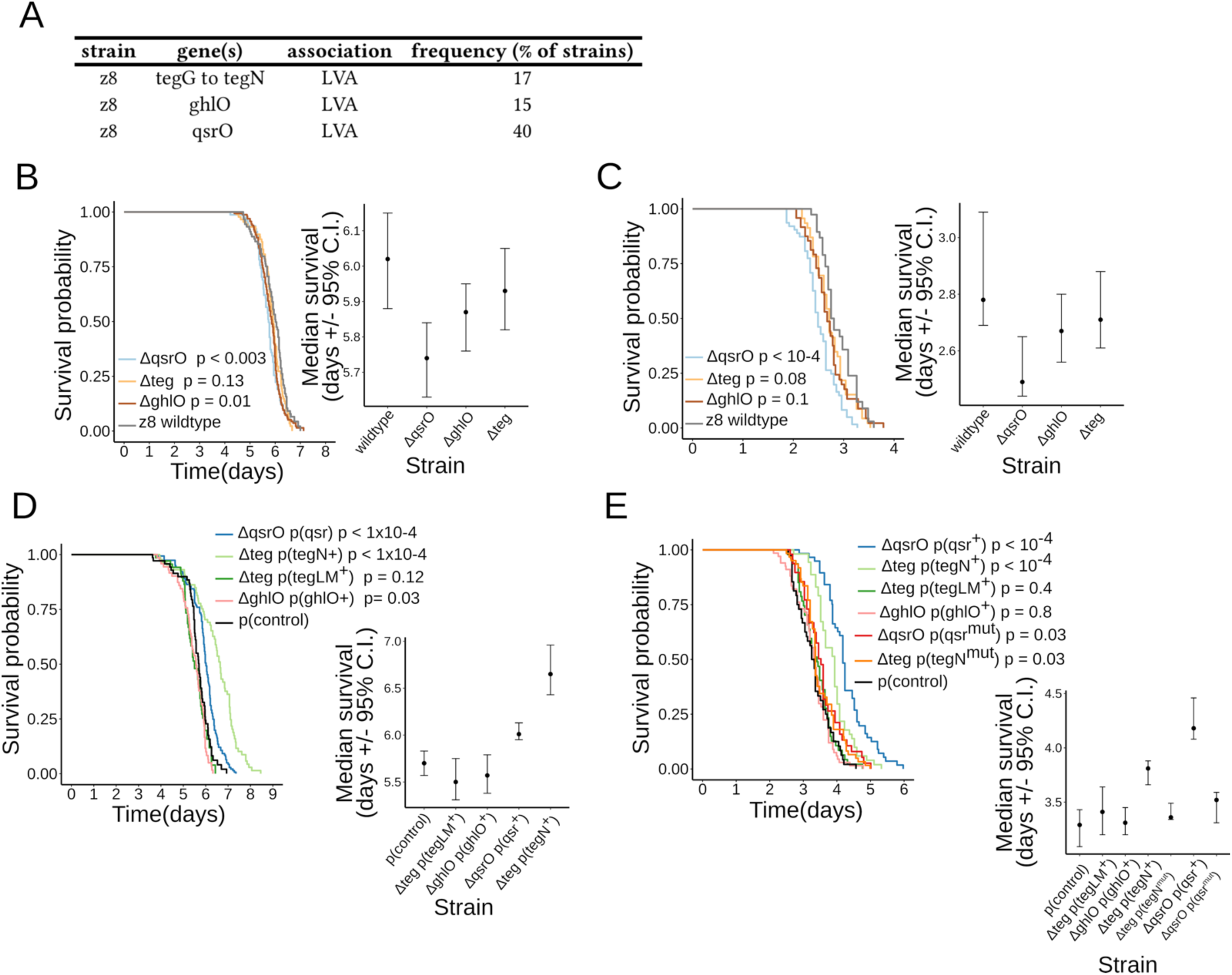
Bacterial virulence upon loss or expression of genes associated with lower virulence. **A)** Summary of the tested LVA genes. Strain, gene nomenclature and gene frequency are indicated. **B-C**) Survival curves and median survival (confidence interval, C.I.) of two strains of adult *C. elegans* worms exposed to three strains of *P. aeruginosa* z8 with deletions in genes associated with lower virulence (*i.e. ΔqsrO; Δteg; ΔghlO*). Wildtype worms are analyzed in (**B**), *pmk-1(lf)* worms in (**C**). Pairwise comparisons of the survival curves between each strain and the z8 wildtype isolate were done using the logrank test. The test p-values are indicated next to each mutant strain in the legend. **D**) Survival curves and median survival (confidence interval ‘C.I.’) of wildtype adult *C. elegans* worms exposed to four strains of *P. aeruginosa* z8 with plasmids expressing genes in gene blocks associated with lower virulence (*i.e. ΔqsrO* p(qsr^+^); *Δteg* p(tegN^+^); *Δteg* p(tegLM^+^); *ΔghlO* p(ghlO^+^)). Pairwise comparisons of the survival curves between each strain and the z8 wildtype strain with control plasmid (p(control)) were done using the logrank test. The test p-values are indicated next to the corresponding strain in the legend. **E**) Survival curves and median survival (confidence interval ‘C.I.’) of *pmk-1(lf)* adult *C. elegans* worms exposed to six strains of *P. aeruginosa* z8 with plasmids expressing genes associated with lower virulence. Four bacterial strains express wildtype bacterial genes (*i.e. ΔqsrO* p(qsr^+^); *Δteg* p(tegN^+^); *Δteg* p(tegLM^+^); *ΔghlO* p(ghlO^+^)). Two additional bacterial strains express mutated bacterial genes (*i.e.ΔqsrO* p(qsr^mut^); *Δteg* p(tegN^mut^)). Pairwise comparisons of the survival curves between each strain and the z8 wildtype strain with control plasmid (p(control)) were done using the logrank test. The test p-values are indicated next to the corresponding strain in the legend.

We constructed strain z8 mutants carrying in-frame deletions of *ghlO, qsrO* and the *teg* gene block (*ΔghlO, ΔqsrO* and *Δteg*, respectively, see also Supplemental Table 5) and measured virulence on two *C. elegans* strains: wildtype and *pmk-1(lf)* mutant. The *pmk-1(lf)* mutant has an impaired p38/PMK-1 pathway that compromises the worm’s response to *P. aeruginosa* PA14 (Kim et al., 2002) and z8 strains (Figure 5B-C). This worm mutant was used as a strain with a genetically ‘sensitized’ background. Deletion of *ghlO* led to marginally reduced survival of wildtype worms (Figure 5B) but not of *pmk-1(lf)* worms (Figure 5C). Deletion of *qsrO*, but not of *teg*, led to a significant reduction in the survival of wildtype worms, indicating an increased virulence of the *ΔqsrO z8* bacteria (Figure 5B). Similarly, deletion of *qsrO*, but not of *teg*, led to a mild but significant reduction in the survival of *pmk-1(lf)* worms (Figure 5C). These results support a direct negative role for the *qsrO* gene in the regulation of virulence. Interestingly, the *qsrO* gene had been reported previously to have a negative regulatory function on quorum sensing (QS), a key contributor to *P. aeruginosa* virulence (Köhler et al., 2014).

To test if the selected genes associated with low virulence can modulate virulence when their expression is enhanced, we constructed strains containing multi-copy plasmids that encode the *ghlO* gene (p(ghlO^+^)), the *qsr* operon (p(qsr^+^)), and *teg* block genes (p(tegLM^+^) and p(tegN^+^)) driven by their native promoters in their respective mutant backgrounds (Supplemental Table 5). The virulence of these strains was measured and compared to a strain carrying an empty plasmid control (p(control)). The virulence of strains overexpressing the *qsrO* and *tegN* genes was significantly reduced compared to the control (Figure 5D, p-value < 10^−4^). In contrast, no difference in virulence was observed for strains overexpressing the *qsrO* and *tegLM* genes (Figure 5D, p-value > 0.01). Strains overexpressing *qsrO* or *tegN* also displayed reduced virulence when tested on immunocompromised *pmk-1(lf)* (Figure 5E, p-value < 0.01). This effect of diminished virulence was abolished when the *qsrO* and *tegN* genes in the plasmids were mutated by introduction of an early stop codon (p(qsr^mut^) and p(tegN^mut^), Figure 5E, p-values > 0.01, see also Supplemental Table 5).

These results suggest a direct role for the *qsrO* and *tegN* genes in the negative regulation of virulence. By contrast, our results suggest the associations of *mexZ, ghlO*, and *tegL* and *tegM* genes with high virulence may not reflect direct causal roles in virulence *per se*. Rather, these latter associations may be secondary to additional underlying factors related to physiological or ecological linkages to virulence. In light of these findings that at least some genes of the accessory genome of *P. aeruginosa* (for example, *qsrO* and *tegN*) can directly modulate virulence implies that processes of selective gene deletion and acquisition (such as horizontal gene transfer, HGT) are critical for the evolution of *P. aeruginosa* virulence in the wild. In summary, the present gene association study identifies 4 previously characterized virulence genes (i.e. PA14_27700, PA14_27690, *pslM, pslK*). In addition, we genetically tested 11 LVA genes by deletion approach, and 6 of these LVA genes by an expression approach, identifying direct roles for *qsrO* and *tegN* in reducing virulence. Importantly, *tegN* is evolutionarily gained or lost altogether with a defined set of 8 accompanying neighboring *teg* genes, i.e. in a physically linked ‘gene block’ (see below, and Supplemental Table 2). Thus, all *teg* genes show association with virulence by being linked to a bona-fide virulence modifier gene (i.e. *tegN*), even though some may not have direct effects on virulence (e.g. *tegM*). A similar pattern is found in other associated genes, that are also found in physically linked gene blocks, and are evolutionarily gained or lost as units (e.g. *qsrO*, PA14_27700).

### The *teg* block is a mobile genetic element that impinges on virulence

Our gene association analysis revealed the *teg* genes (i.e. genes *tegG* to *tegN*), as a set of LVA genes. Among the experimental isolate collection, strains where this group of *teg* genes is present had lower virulence compared to those where it is absent (Welch t-test, *p*-value = 0.005), as expected from the gene association results. Our finding that *tegN* directly modulates virulence when expressed (Figure 5D-E), strongly suggests a functional link between the *teg* genes and reduced virulence.

To better understand the organization of the *teg* genes and their possible mode of acquisition/loss, we examined features of the *tegN* locus by *in silico* analysis of three *P. aeruginosa* isolates with complete genomes (strains atcc27853, SCV20265 and PA7790) that allow uninterrupted examination of chromosomal features and synteny around *tegN*. The *teg* locus contains a conserved genomic repeat of ∼7 kilobases (Figure 6A). This genomic repeat is found in 2-4 tandem copies in the queried genomes (Figure 6A). The repeats are not completely identical between strains, and display stretches of varying conservation (Figure 6B). We refer to this tandem genomic repeat unit as the ‘*teg* block’.

**Figure 6.**
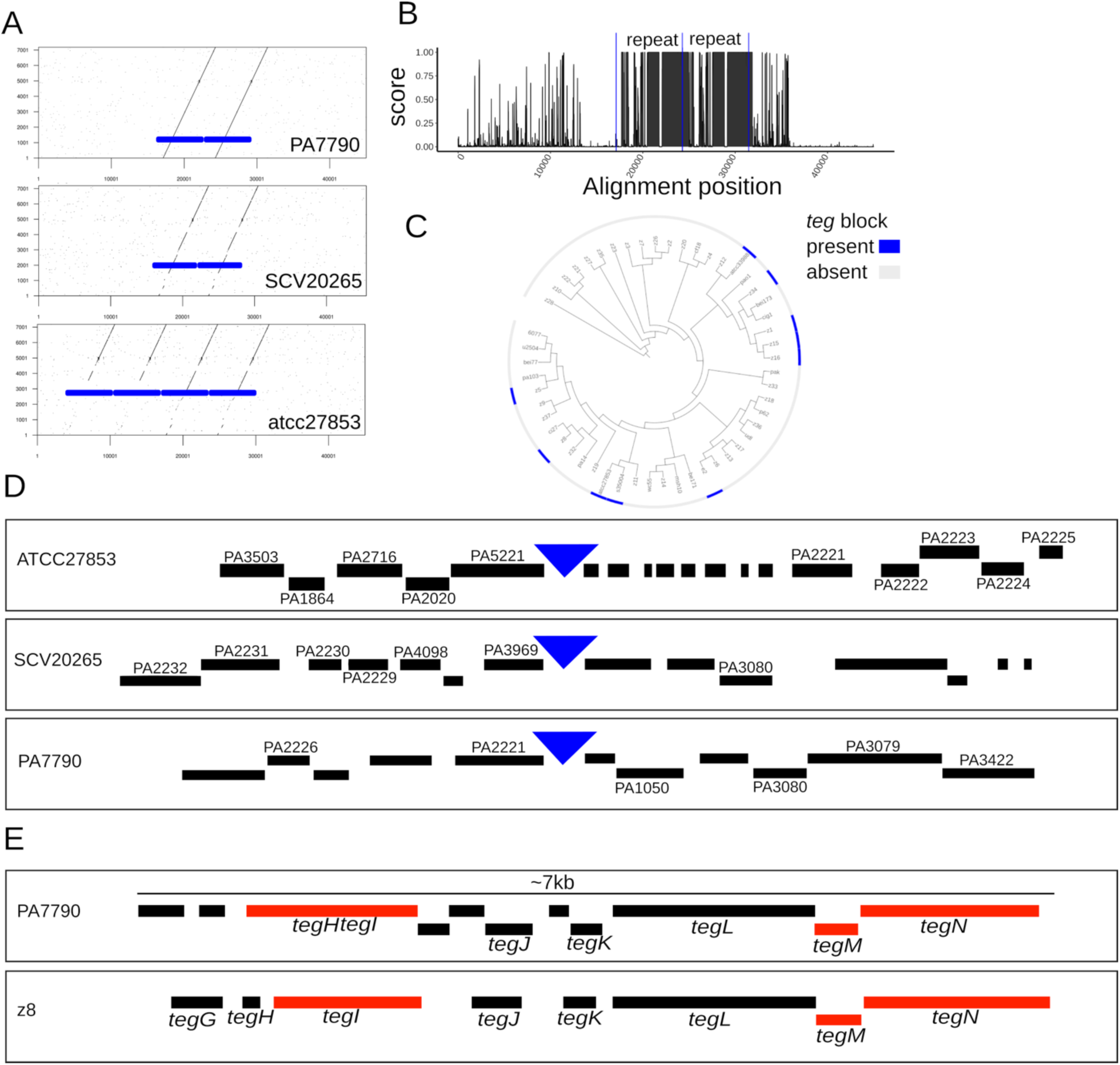
Features of the LVA-associated *teg* block. **(A)** Dot plot comparison between the *teg* block repeat from strain PA7790 (y-axis) and homologous genomic regions in 3 strains with complete genomes (PA7790, SCV20265 and atcc27853). Solid blue boxes indicate the tandem repeat sequence observed. (**B**) Sequence conservation (PhastCons score) for the alignment of the *teg* block genomic regions displayed on (A). The presence of two tandem repeated regions is indicated between the vertical blue lines. (**C**) Phylogenetic distribution of the *teg* block in the 52-strains isolate set. Presence (blue color) or absence (grey color) of the *teg* block is indicated. The block is found in 10 strains in different clades. (**D**) Gene neighborhood around the *teg* block insertions sites (blue triangles) in strains atcc27853, SCV20265 and PA7790. Predicted protein-coding genes are indicated by black boxes. Genes with homologs in the PAO1 strain are named. (**E**) Gene presence in the *teg* block repeat of strains PA7790 and z8. Predicted protein-coding genes are indicated by black and red boxes. Red boxes indicate genes with links to viral-related function. Eight genes in *teg* block of strain z8 are named (*tegG* to *tegN*).

The frequency and phylogenetic distribution of the *teg* block in the 52-strain collection suggest that the element is mobile. The block is found in 10 strains, corresponding to 19% of the collection (Supplemental Table 1), and it is distributed to multiple clades (Figure 6C). The simplest hypothesis to account for the phylogenetic pattern of the *teg* block is seven independent acquisitions. A comparison of the genomic neighborhoods surrounding the location of the *teg* block in the 3 complete genomes showed no evident synteny (Figure 6D), arguing against an ancestrally fixed genomic location, and also supporting the conclusion that the *teg* block is a mobile genetic element. Curiously, two genes (PA2221, PA3080) were commonly shared in 2 distinct pairs of neighborhoods.

The predicted proteins encoded by the *teg* block also supports genetic mobility as a potential function. The conserved repeat unit (i.e. *teg* block) has 8 and 11 predicted protein-coding genes in strains PA7790 and z8, respectively, and includes the *tegG* to *tegN* set, named and investigated in strain z8 (Figure 6E). Five of the predicted *teg* proteins (*tegG, tegH, tegJ, tegK, tegL*) have no features or annotations that could help infer their functions. However, three of the *teg* proteins have features and annotations that suggest virus-related functions. The gene *tegI* encodes a viral ‘replication initiation protein’ homologous to *gpII* of phage M13. *tegM* encodes a homologue of viral coat protein g6p of phage Pf3, with a conserved DUF2523 domain (CDD domain accession: pfam10734). *tegN* encodes a P-loop containing NTPase (CDD domain accession: cl21455), a homologue of *gpI* found in phage M13. These annotations suggest that the *teg* block encodes functions related to DNA replication (*tegI*) and virion assembly (*tegM* and *tegN*) (Loh et al., 2017; Ledsgaard et al., 2018), supporting the conclusion that the *teg* block is a virus-related element. The apparent absence of proteins with functionality for chromosomal integration or conjugative transfer may indicate that the *teg* block may rely on proteins from its bacterial host or other mobile genomic elements for these putative functions.

### Genomic presence of the *teg* block is restricted by CRISPR-Cas systems

The composition of the *P. aeruginosa* accessory genome is shaped by uptake of genes from other microorganisms via horizontal gene transfer (HGT), frequently involving mobile genetic elements (MGE) such as prophages and ICEs (integrative and conjugative elements). HGT events can be restricted by diverse classes of bacterial defense systems, which protect cells against the acquisition of elements that could confer deleterious phenotypes. Since we observed that the *teg* block, a viral-like element of the *P. aeruginosa* accessory genome, associates and negatively regulates virulence, we investigated if such element would be restricted by the bacteria.

We first explored the possibility that CRISPR-Cas systems could restrict the uptake of the *teg* block. For this purpose, we utilized the existence of an immunity record in the CRISPR spacer loci of *P. aeruginosa* strains. CRISPR repeat spacer sequences identify genes whose restriction by CRISPR-Cas systems of *P. aeruginosa* has been selected for during the recent evolution of the strains examined. Except in rare cases of apparent spacer ‘self-targeting’ (Stern et al. 2010; see below), CRISPR spacers and their protospacer target genes are predominantly found in different genomes.

We identified the set of all CRISPR spacers present in 1488 strains and searched for their targets in the *P. aeruginosa* pangenome. In this manner, we identified 693 genes that are targeted by spacers (Supplemental Table 6). The vast majority (670 out of 693, corresponding to 97%) of the identified spacer-targeted genes are not found on the same genomes as the spacers that target them, and thus reflect genes whose integration into the genome of a given strain was successfully blocked by CRISPR-Cas during the evolution of that strain. We next determined the relationship of the spacer-targeted genes to virulence. At the single gene level, the vast majority of the spacer-targeted genes (683) showed no statistically significant correlation with virulence (Figure 7A-B). Nonetheless, a set of 9 genes was associated with low virulence (*i.e*. LVA genes, Figure 7B, p-value < 0.01 by M-W test). In contrast, only one spacer-targeted gene (cluster #18193) showed significant association with high virulence.

**Figure 7.**
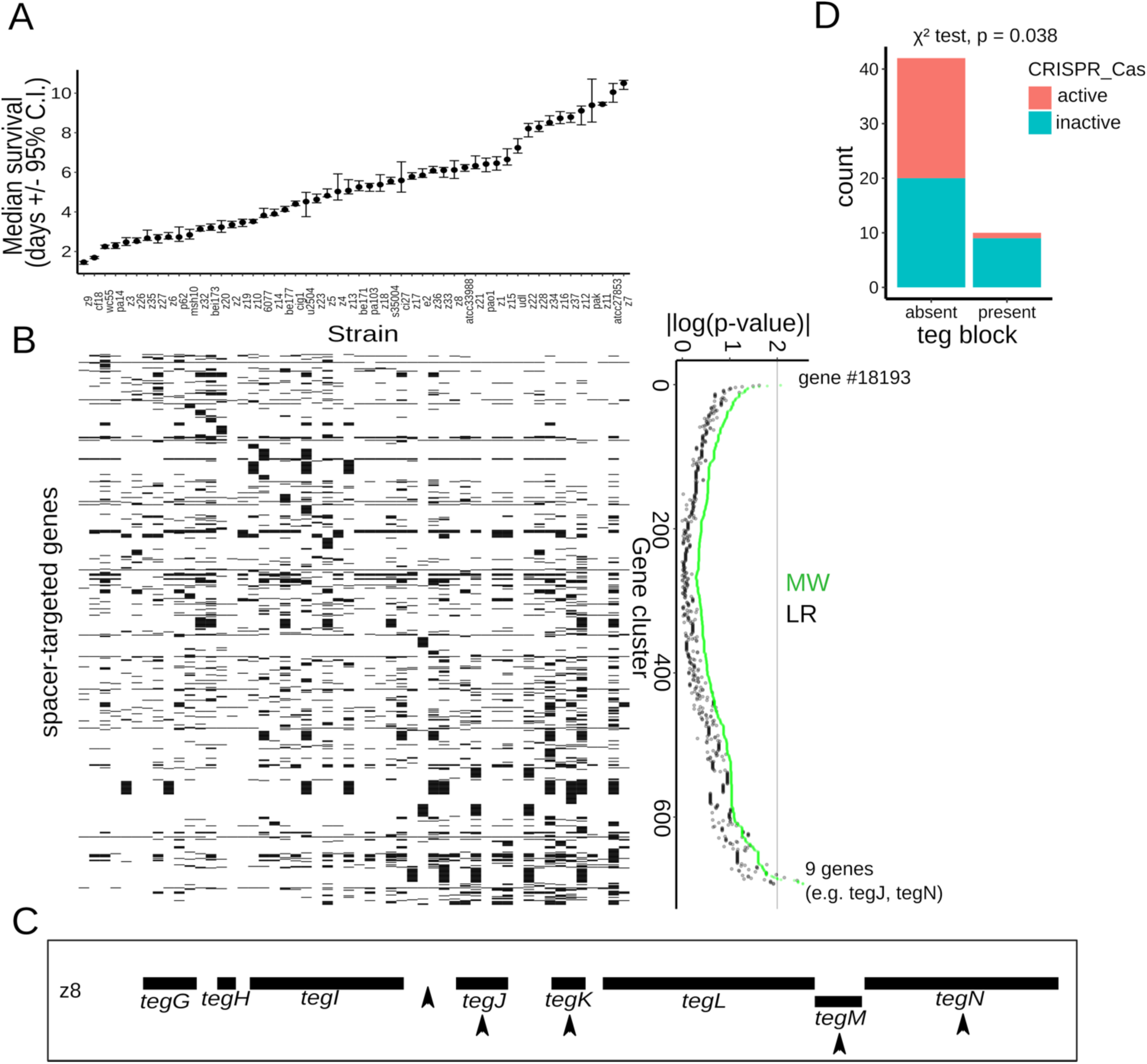
Restriction of the *teg* block by spacers and correlation with CRISPR-Cas systems. (**A**) Median survival of adult *C. elegans* worms exposed to the studied collection of *P. aeruginosa* strains. The strains are ordered from high to low virulence (left to right) and aligned with the matrix below. (**B**) Left panel: gene presence/absence matrix for genes targeted by CRISPR spacers. Gene presence is indicated with black squares and absence with white squares. Genes (rows) are aligned with the corresponding p-values. Right panel: Association statistics (p-value of MW and LR tests) for the CRISPR-targeted genes, shown as | log_10_(pval)|. Rows are ordered from association with high virulence to association with low virulence. (**C**) Schematic of the *teg* block in strain z8. Black boxes indicate *teg* genes and arrowheads spacers that target the element. (**D**) Number of strains (i.e. count) where the *teg* block is present or absent in relationship with the status of the host CRISPR-Cas system (active in red color, inactive in cyan color). The p-value of a Chi square test is indicated.

Among the LVA spacer-targeted gene set, 5 out of 9 genes were found to be genes in the *teg* block (Figure 7C). Thus, the spacer-encoded immunity record shows repeated restriction of the *teg* block by CRISPR-Cas systems, consistent with it being detrimental to bacteria. Additional spacer-targeted genes, included mostly genes of unknown function, although some annotations related them to mobile elements (*i.e*. integrase for gene #6157, ‘phage capsid’ for gene #8274) as expected.

Considering that the spacer-encoded record of restricted genes is finite and reflects recent restriction events, we evaluated the *teg* block presence or absence in relationship to the genomic presence or absence of CRISPR-Cas systems in the isolates. Significantly, the ‘*teg* block’ is found predominantly among strains with inactive/absent CRISPR-Cas systems (9/10 strains, Fig 7D, Welch t-test, p-value = 0.038). Altogether, these results show that the *teg* block, a virulence-inhibiting viral-like accessory genome element, is restricted by CRISPR-Cas systems, as indicated by the pangenomic presence of spacers targeting it, and its predominant presence in strains without active CRISPR-Cas systems.

### Active CRISPR-Cas systems positively but indirectly correlate with *P. aeruginosa* virulence

Extending our analysis beyond the *teg* block, we analyzed the overall statistical features of the spacer-targeted genes. The statistical distribution of the gene association statistic (p-value of the LR test) revealed that the set of spacer-targeted genes, associates preferentially with lower virulence, when compared to not spacer-targeted genes (Fig 8A, two sample K-S test, p-value 7×10^−12^). Furthermore, the statistical distribution of spacer-targeted genes separated by their affiliation to higher or lower virulence also differs significantly (Fig 8B, two sample K-S test, p-value 2.2×10^−16^), and this difference in the distributions remains upon removal of the *teg* loci from the comparison (two sample K-S test, p-value 2.2×10^−16^). Altogether, these results suggest that spacer-targeted genes are enriched in their association with lower virulence, and this enrichment is driven by a plethora of gene associations, in addition to those of the *teg* genes. Moreover, we anticipate that association studies using larger isolate collections should allow better resolution of the individual gene association scores, and may assist in identification of additional spacer-targeted LVA genes.

**Figure 8.**
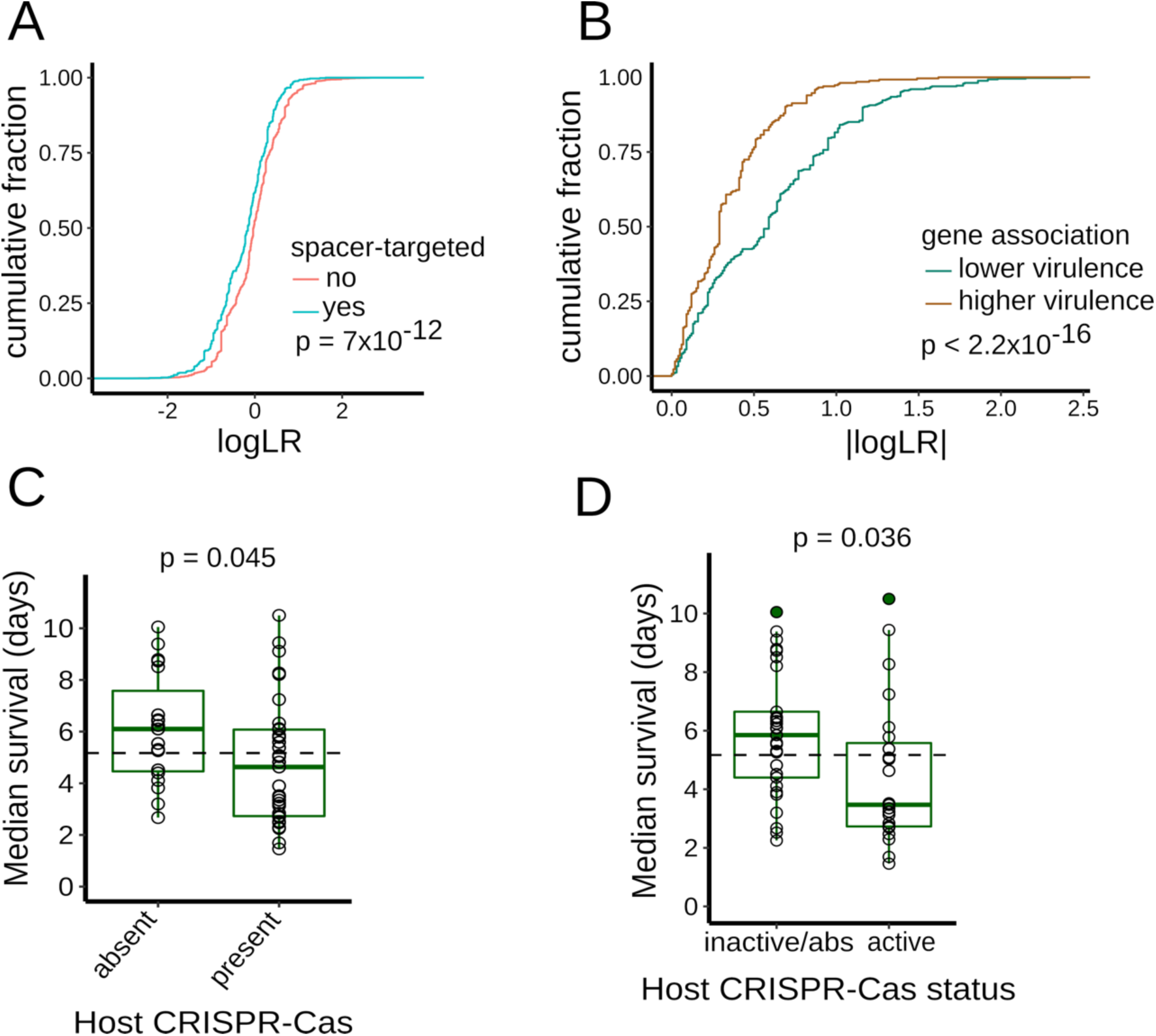
Relationship between virulence and spacer-targeted genes and CRISPR-Cas systems of *P. aeruginosa*. (**A**) Cumulative distributions for the association statistic (log p-value of LR test). Genes in the accessory genome are partitioned according to the whether they are targeted by spacers (in cyan color) or not (in red color). The p-value of two sample K-S test is indicated. (**B**) Cumulative distributions for the association statistic (log p-value of LR test). Spacer-targeted genes are partitioned according to the whether they are associated with higher (in brown color) or lower (in green color) virulence. The p-value of two sample K-S test is indicated. (**C-D)** Box plots of worm median survival in relationship with CRISPR-Cas presence/absence and activity status. **C**) Strains are partitioned according to the presence/absence of host CRISPR-Cas systems (I-E, I-F). **D**) Strains are displayed according to their CRISPR-Cas status in two categories: active, or inactive-absent (inactive/absent). The median virulence of the complete set of strains displayed on each graph is indicated with the dashed horizontal line. p-values are indicated for the Welch *t*-test comparison of virulence between the two groups represented (C-D).

Since we observed that elements of the *P. aeruginosa* accessory genome can negatively associate with virulence, and such elements can be restricted by bacterial CRISPR-Cas systems, we used gene association analysis to test for the association of virulence against *C. elegans* with the presence or absence of restriction-modification (RM) systems, CRISPR-Cas systems, and a recently identified cohort of ten novel defense systems (Doron et al., 2018). These kinds of defense systems are widely distributed in bacteria and display innate (RM systems) or adaptive immune characteristics (CRISPR-Cas systems). We first analyzed adaptive immune systems on the premise that these systems may be able to selectively filter out deleterious genetic elements.

Type I CRISPR-Cas systems (Cas proteins and spacer arrays) are present in 71% of the 52 strains (37/52 strains; Supplemental Table 1) and belong to three different subtypes, that can be absent/present independently of each other: type I-F (73%), type I-E (35%) and I-C (21%). This distribution of CRISPR-Cas systems is consistent and similar to previous surveys of *P. aeruginosa* CRISPR-Cas systems (van Belkum et al., 2015). In addition to the genomic presence of CRISPR-Cas loci, we also investigated if the identified CRISPR-Cas systems were predicted to be active or inactive based on the presence/absence of known anti-CRISPR genes. Anti-CRISPR proteins, are virus-encoded and can inhibit CRISPR-Cas systems, blocking their immune function (reviewed in (Pawluk et al., 2017)). We identified a set of 22 anti-CRISPR gene families in 31% of the 52 *P. aeruginosa* genomes and cataloged each strain’s CRISPR-Cas status as: 1) ‘active’ if it has at least one CRISPR-Cas system with no known cognate anti-CRISPR gene present in genome; or 2) having an ‘inactive/absent’ system if CRISPR-Cas is absent or where cognate anti-CRISPR gene(s) are found concomitantly with CRISPR-Cas (Supplemental Table 1).

We compared the above anti-CRISPR approach for identifying strains with inactive CRISPR/Cas to an alternative criterion: the presence in the same bacterial genome, of a CRISPR-Cas spacer with its DNA target, a condition referred to as spacer ‘self-targeting’ (Stern et al. 2010). The presence in the same genome of a CRISPR-Cas locus and one or more self-targeting spacers is considered to reflect an inactive effector status of that CRISPR-Cas locus, because genome cleavage by an active CRISPR-Cas system is expected to be lethal to the bacterial cell (Bikard et al., 2012; Vercoe et al., 2013). In our collection, we found 11 strains with CRISPR-Cas and at least one self-targeting spacer with a full match to its genomic target (Supplemental Table 1). Most of these strains (9 out of 11, corresponding to 82% of them) were included in the set of inactive strains by the anti-CRISPR approach. The determination of CRISPR-Cas ‘inactivity’ with the two approaches is highly similar (McNemar’s Chi-squared test, p-value = 1).

Next, we analyzed the CRISPR-Cas systems in relationship to virulence. We first considered separately the subtypes I-F, I-E, I-C and their combinations (Supplemental Figure 5A). Strains with type I-C CRISPR-Cas systems showed lower virulence compared to that of all other strains (Welch t-test, p-value = 0.03). The distinct association observed for I-C systems coincides with the fact that *P. aeruginosa* type I-C CRISPR-Cas systems have been exclusively found inside pKLC102-like ICEs (van Belkum et al., 2015). Defense systems inside ICEs, such as type I-C CRISPR-Cas systems, likely fulfill a role in the ICE’s lifecycle and may not primarily provide immune protection to the bacterial host. Based on this evidence, we did not consider I-C systems part of *P. aeruginosa* complement of immune systems, and so in subsequent analysis we considered only subtypes I-E and I-F as comprising the bacterial cell’s CRISPR-Cas systems.

Interestingly, we found that the presence of a host CRISPR-Cas system (*i.e*. either subtypes I-E or I-F), significantly associates with higher virulence (Figure 8C, Welch t-test, p = 0.045). To investigate if this association is related to the immune function of CRISPR-Cas systems, we considered the status of activity of the host CRISPR-Cas systems. Notably, the presence of active CRISPR-Cas systems (by the criterion of absence of anti-CRISPR genes) also statistically correlates with increased virulence (Figure 8D, two-sided Welch t-test, p = 0.036). Moreover, upon inclusion of strains with spacer self-targeting to the ‘inactive’ strain set, the statistical association between active CRISPR-Cas and higher virulence is maintained (one-sided Welch t-test, p = 0.038). To further investigate the relationship between CRISPR-Cas and virulence, we applied an alternative analysis. The survival curves for the strain collection were pooled, forming two groups based on the presence or absence of CRISPR-Cas in the isolates. The survival curves between these two groups differ significantly (Supplemental Figure 5B, K-M method, logrank test, p-value < 2×10^−16^), and the strain group with CRISPR-Cas systems has a lower median survival (4.2 days, 95% C.I.: 4.0 – 4.4 days) compared to the group without this defense system (median survival of 6.5 days, 95% C.I.: 6.3 – 6.6 days).

The association of active CRISPR-Cas systems with high virulence suggested a positive role for this immune system in the maintenance of virulence. Thus, we explored whether or not CRISPR-Cas could have a direct role in virulence. First, we constructed a deletion of the entire six Cas genes of strain PA14 (strain PA14ΔCas) to abolish CRISPR-Cas activity, but we observed no significant difference in virulence between the PA14ΔCas and wildtype PA14 (Supplemental Figure 5C). In addition, we tested if the Cas proteins have the ability to modulate virulence when expressed from a plasmid in strain PAO1 that lacks CRISPR-Cas. The PAO1 strain expressing CRISPR/Cas from a plasmid, (strain PAO1 p(Cas^+^), displayed no significant difference in virulence compared to PAO1 expressing a plasmid control (p(control)) (Supplemental Figure 5D). In summary, these results indicate that CRISPR-Cas is neither necessary nor sufficient to directly modulate bacterial virulence, at least under the assayed laboratory conditions.

We next proceeded to analyze known and presumed innate immune systems of *P. aeruginosa*: RM systems (Roberts et al., 2015) and the cohort of ten novel defense systems (Doron et al., 2018), respectively. We identified RM systems based on annotations from the REBASE database (Roberts et al., 2015) (Supplemental Table 1). One or more predicted RM systems are present in 96% of the strains (50/52 strains), with an average of 3.8 RM systems per strain. We observed a weak association between the total number of RM systems and virulence (Supplemental Figure 6A, spearman rank correlation, □: 0.25) that does not reach significance (p = 0.08). Similarly, the relationship between each separate RM system type and virulence shows weak association for the types I, II, III and no association for type IV RM systems (Supplemental Figure 6). None of the above-mentioned correlations reached statistical significance (all p-values >= 0.08). Next, we evaluated the presence of ten novel defense systems (Doron et al., 2018) by homology of the system’s diagnostic proteins to genes in our strain collection. We identified most of the novel defense systems (8 out of 10) identified by Doron et al. (2018) in the 52 strains set (Supplemental Table 1). The lammassu, septu, zorya and hachiman systems were found in a low number of strains (2-8% frequency). In contrast, the druantia, shedu, wadjet and gabija systems occurred at higher incidence in *P. aeruginosa* (14-37% frequencies). We found no statistically significant association with virulence for any of the novel immune systems (Supplemental Figure 7. Similarly, we observed no association between the overall number of novel defense systems per strain and virulence (spearman rank correlation, r: 0.03, p = 0.81, Supplemental Figure 7). These results show that the presence or absence of the recently identified immune systems bears no apparent relationship with strain virulence. Interestingly, we noted that the gabija system of strain PA14 (genes PA14_60070 and PA14_60080) and strain CF18 (genes #2421 and ID #Q002_01766) are found inside ICEs: PAPI-1 (He et al., 2004) for PA14, and an unnamed ICE (predicted with ICEfinder (Liu et al., 2019)) for CF18. Altogether, these observations highlight that ICEs can harbor multiple defense systems, as previously exemplified with type I-C CRISPR-Cas systems.

In summary, we found that RM and novel defense systems have a weak or no significant relationship with virulence. In contrast, the presence and activity of CRISPR-Cas systems associates with higher virulence. The statistical association between active CRISPR-Cas systems and *P. aeruginosa* virulence suggests that CRISPR-Cas activity may indirectly affect virulence-related phenotypes, most likely by regulating acquisition and/or retention of accessory genome virulence factors and other elements that impinge on virulence. A verified instance of such CRISPR-Cas mediated restriction process, is exemplified by the *teg* block. Moreover, the statistical distribution of the gene association statistic for the spacer-targeted genes suggest the possibility that additional restricted LVA genes may be identified in more powerful association studies.

## DISCUSSION

In the present study, we investigated bacterial-driven variation in the interactions between *C. elegans* and *P. aeruginosa*. 52 *P. aeruginosa* wild isolate strains were found to cover a wide virulence range, spanning from highly virulent strains, which induce a worm median survival of 1.5 days (∼11% of their lifespan under standard conditions at 25°C) to strains with almost no virulence, which induce worm lifetimes similar to those observed with non-pathogenic *E. coli* HB101, and which do not affect progeny production.

Considering that *P. aeruginosa* is a free-living bacterial species that facultatively engages in pathogenic interactions with invertebrates, and considering that *C. elegans* is a natural bacterial predator, it is conceivable that the strain variation in virulence towards *C. elegans* reflects adaptations of *P. aeruginosa* to its natural niches. In natural settings virulence may be a character under selection by the frequency with which predators are deterred by virulence mechanisms, and/or by the extent to which the bacterium depends on infection of predator hosts for population growth.

It should be noted that because *P. aeruginosa* is a multi-host pathogen of many species, including insects and single-celled eukaryotes, as well as nematodes, we cannot say with any certainty whether any of *P. aeruginosa* strains chosen for this study have undergone selection in the wild through direct interaction with *C. elegans*. We observed that amongst our 52-strain panel, environmental strain isolates exhibited on average greater virulence against *C. elegans* than did clinical isolates (Supplemental figure 1B), consistent with previous findings (Sánchez-Diener et al., 2017). This suggests that some of the strain variation in virulence against *C. elegans* could be influenced by adaptations of *P. aeruginosa* to its pathogenic association with humans, and that such adaptations may not necessarily confer pathogenic benefit against *C. elegans*. The virulence of clinical isolates could reflect genetic and genomic makeup of the bacterium that is favorable in the context of human immune responses and/or therapeutic antibiotics. Indeed, among the genes associated with virulence, we observed several genes involved with antibiotic resistance, such as *mexZ*, a negative regulator of the *mexXY* bacterial efflux pump (Aires et al., 1999; Westbrock-Wadman et al., 1999) and *arr*, which functions to induce biofilms in response to aminoglycoside exposure (Hoffman et al., 2005).

The variation in virulence among *P. aeruginosa* strains parallels the substantial genomic diversity of this bacterial species. *P. aeruginosa* strains contain relatively large genomes for a prokaryote (5-7 Mb; 5000-7000 genes) with a sizable contribution of accessory genome elements (Figure 1). Our data show that strain variation in *P. aeruginosa* virulence is mediated by specific accessory genome elements (Figures 3-4), in combination with the core genome, including previously described *P. aeruginosa* virulence-related factors (Figure 4). Notably, we find particular accessory genome elements that contribute to increased virulence, and others that promote decreased virulence (Figures 3 and 5). The existence of genes whose functions lead to the negative regulation of virulence (for example, *qsrO* and *tegN*) suggests: 1) strain adaptations to niches where capping virulence is advantageous, either for environmental reasons (*e.g*. infrequent bacterial predators or hosts for bacteria to feed on) or clinical reasons (*e.g.* evasion of immune surveillance at lower virulence); 2) detrimental effects of MGEs (e.g. *teg* block) that are chromosome integrated and likely engage into parasitic relationship with its bacterial host.

The results of our genetic analysis of HVA and LVA genes indicate a direct role for a subset of these genes in modulating virulence, whereas for other HVA and LVA genes our genetic results do not support a direct role. A direct role in virulence for genes PA14_27700, PA14_27680, *pslK*, and *pslM* was expected based on previous findings (Figure 4) and hence their identification as HVA genes supports our comparative genomics approach. For 11 LVA genes that we tested genetically, the results suggest a direct contribution for *qsrO* and *tegN* to virulence (Figure 5). On the other hand, genetic ablation (for t*egG* to *tegN* and *ghlO*) or ectopic expression of *mexZ, tegL, tegM, ghlO* (Figure 5, Supplemental Figure 4) or the *Cas* genes (Figure 7) did not measurably alter virulence. Importantly, associated genes can be evolutionarily gained or lost as multigene units -- physical blocks with defined sets of accompanying neighboring genes. Genes in such blocks all show association with virulence by being linked to a bona-fide virulence modifier gene, even though some may not have direct effects on virulence. This situation is exemplified by the *teg* block, that comprises 8 LVA genes (Fig. 6), including one that affects virulence (i.e. *tegN*) and others that do not (i.e. *tegL, tegM*).

What could account for why certain genes would not exhibit essential virulence functions in genetic tests, despite being correlated with virulence in gene association analysis? One possibility could be statistical false discoveries. However, we assessed the reliability of our statistical analysis in two ways: by using permutation-based testing to filter out false discoveries, and by employing phylogenetically aware scoring approaches to control for any confounding effect mediated by population structure.

It is also possible that some of the genes that tested negatively in the genetic tests actually do function in some contexts as *bona fide* virulence factors, but their effects could be masked by epistasis in the genomic background of the particular strains in which we conducted our loss-of-function and gain-of-function tests. The possibility of such strain-specific epistasis could be investigated by conducting parallel genetic tests for the full cohort of relevant strains.

Our analysis of the *teg* block illustrates that LVA genes can reside within MGEs that are detrimental for virulence (Figure 5) and that are restricted by host CRISPR-Cas systems (Figures 6 and 7). The *teg* block is likely not the only MGE with a negative association to virulence, because the cohort of spacer-targeted genes shows an overall enriched association with lower virulence (Figure 8A-B). We thus suggest that additional MGEs, detrimental for virulence and CRISPR-Cas restricted, could be unveiled utilizing more powerful association studies with enlarged isolate collections.

We observe a positive correlation between the virulence of *P. aeruginosa* strains against *C. elegans* and the presence of CRISPR-Cas bacterial immunity (Figure 8C-D), even though our genetic tests with CRISPR-Cas loss-of-function mutants or ectopic expression indicate that CRISPR-Cas activity is neither necessary nor sufficient for increased virulence (Supplemental figure 5C-D). This suggests that bacterial adaptive immunity and anti-predator virulence may be somehow indirectly coupled via the effects of physiological, ecological, and/or evolutionary factors.

Although there are undoubtedly numerous potential underlying causes for a linkage between CRISPR-Cas and virulence, two broad classes of potential scenarios are suggested (Figure 9). One scenario is based on possibility that the evolution of accessory genomes is highly influenced by bacterial restriction systems, such as CRISPR-Cas that function to limit horizontal gene transfer (HGT) and thereby help shape the makeup of the accessory genome. Our finding that accessory genome elements can modulate virulence supports the supposition that bacterial immune systems could indirectly contribute to the maintenance or evolvability of virulence towards invertebrate predators such as *C. elegans.* This scenario is further supported by our findings that *P. aeruginosa* genes associated with low virulence include detrimental viral-like mobile genetic elements and are more enriched for targeting by CRISPR-Cas spacers that are those associated with higher virulence.

**Figure 9.**
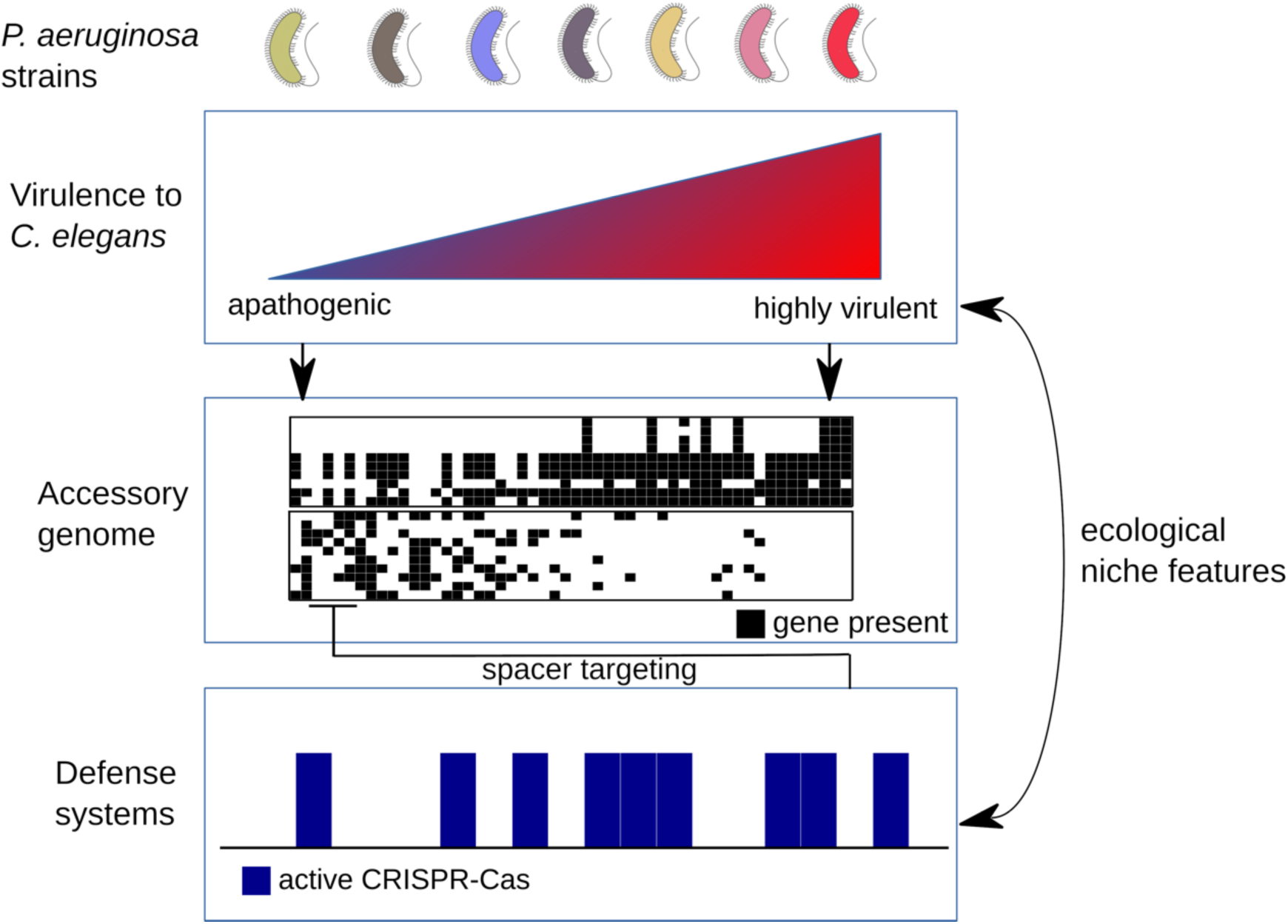
Working Model for linkage between CRISPR-Cas and virulence. The strain diversity of *P. aeruginosa* encompasses an ample range of virulence towards adult *C. elegans* worms. Accessory genome elements, represented by a matrix of gene presence/absence (in black/white boxes, respectively), correlate and contribute to the differences in virulence among strains (indicated by arrows). Active defense systems, such CRISPR-Cas, are enriched in strains with higher virulence. CRISPR spacers target accessory genome elements that are associated with lower virulence (negative arrow). The distribution of active defense systems and higher virulence may also reflect, the co-occurrence of frequent bacterial predators and frequent viral infections in some of the niches inhabited by *P. aeruginosa*.

A second scenario, not mutually exclusive with the first, is based on the fact that bacterial restriction systems such as CRISPR-Cas are themselves often part of the accessory genome, as evidenced in the case of *P. aeruginosa* by the fact that some strains contain one or more CRISPR-Cas loci, while other strains contain none. Apparently, CRISPR-Cas adaptive immunity is selected for or against, depending on particular environmental conditions. Therefore, high virulence and the capacity to restrict HGT could be linked by the co-occurrence of environmental factors that simultaneously select for both features. For example, in certain *P. aeruginosa* natural habitats, abundant predation by invertebrates such as *C. elegans* may commonly co-occur with pressure from an abundance of phages. Conversely, clinical settings may frequently present conditions that simultaneously disfavor high virulence and restriction of HGT. Testing of these hypotheses will benefit from further studies.

Unlike active CRIPSR-Cas, we did not observe a similar association of virulence with other restriction systems, including restriction/modification (RM) and a set of recently identified restriction systems of less well characterized mechanisms (Doron et al., 2018). These other systems, particularly the RM systems, differ from CRISPR-Cas fundamentally in that they are not adaptive immune systems, and hence they would tend to limit uptake of foreign DNA elements regardless of whether those elements confer positive or negative phenotypes. CRISPR-Cas systems are much more discriminatory: Restriction of an element by CRIPSR-Cas requires programming the spacer array with a sequence from the targeted element, enabling selection for targeting of deleterious elements, and selection against targeting of advantageous elements. Thus, the association that we observe between virulence and active CRISPR-Cas may reflect such selection for restriction of uptake of elements that are particularly deleterious in the context of anti-predator virulence.

This study shows that genome-wide association (GWAS) analysis of a panel of genomically-diverse strains of a bacterial species can identify previously-unrecognized accessory genome elements influencing a phenotype of interest, in this case virulence of *P. aeruginosa* against the invertebrate bacterivore *C. elegans*. What sorts of genetic bases for virulence variation might have been missed in our study? First, some of the accessory genome genes that scored below statistical cutoffs in our study might emerge as high-confidence candidate virulence modulators from studies of larger and/or more diverse panels of bacterial strains. It should also be noted that our gene-association analysis scored for presence or absence of intact (accessory genome) genes. We did not attempt to test for association of virulence with amino acid coding mutations, or with noncoding sequence polymorphisms that could alter *cis-*regulatory regulation of direct virulence modulators. Such higher resolution (GWAS) analysis could be the basis for future inquiries.

## MATERIALS AND METHODS

### *C. elegans* worm strains

The *C. elegans* N2 strain was used as wild type strain. In addition, strain KU25: *pmk-1(ku25)*, referred to as *pmk-1(lf)*, was used for some virulence assays. All nematode strains were maintained using standard methods on NGM plates (Brenner, 1974) and fed with *E. coli* HB101.

### Bacterial strains

The *P. aeruginosa* strains were routinely grown on LB media at 37°C without antibiotics, unless otherwise noted. A list of the 52 bacterial isolates established as our experimental panel is listed in Supplemental Table 1. The collection was assembled using strains procured from numerous distinct sources, and although we strove to obtain a diverse collection of both environmental and clinical stains, there was limited control over the collection composition with regard to specific features. The description and genotypes of bacterial strains constructed in the present study are listed in Supplemental Table 5. For a portion of the strains in the collection, we found that genetic manipulation is limited, because a considerable fraction of the isolates exhibit strong restriction to uptaked DNA or high levels of resistance to antibiotics.

### Worm survival assays (Virulence assays)

Worm survival assays (virulence assays) were performed using slow killing (SK) conditions (Tan et al., 1999). Briefly, an aliquot of an overnight liquid LB culture of each *P. aeruginosa* strain was plated on SK agar plates. The bacterial lawn was spread to cover the entire surface of the agar, to prevent worms from easily escaping the bacterial lawn. The plates were incubated at 37°C for 24h and then at 25°C for 24h, to allow growth of the lawn and the induction of pathogenic activity (Tan et al., 1999). Prior to use, FUDR (100 ng/µL) was added to the plates to a final concentration in the agar medium of 300 µM. A synchronous population of young adult (YA) hermaphrodite N2 worms was prepared by standard hypochlorite treatment, followed by culture of larvae from L1 stage to YA stage on NGM agar seeded with *E. coli* HB101. The young adult (YA) worms were then transferred to the SK plates to initiate their exposure to *P. aeruginosa* lawns. The time-course of death of the worms on each plate was determined with the aid of a “lifespan machine” (Stroustrup et al., 2013), an automated system based on a modified flatbed scanner. Image analysis was optimized to fit the *P. aeruginosa* slow killing conditions as described previously (McEwan et al., 2016). The collected survival information was manually curated and analyzed using R (i.e. survminer package) with the Kaplan-Meier (K-M) method. K-M was used to estimate median survival and its confidence interval. Survival curves are representative of at least two independent experiments. The K-M based estimate of the ‘median survival’ of worms exposed to a particular bacterial isolate corresponds to our measure of bacterial virulence. The semiparametric Cox proportional hazards model, is not applicable to the obtained survival information, as the proportional-hazards (PH) assumption does not hold (R ‘survival’ package, proportional hazards test, global p-value = 0; p-value < 0.05 for 15 strains).

In the alternative analysis of the survival data to study the relationship of virulence to CRISPR-Cas, the survival data (i.e. individual worm lifespans) of all strains with host CRISPR-Cas systems was aggregated into a first group (n = 2656), and the survival data for strains without host CRISPR-Cas systems was aggregated into a second group (n = 1549). The aggregated data was analyzed using R (i.e. survminer package) with the Kaplan-Meier (K-M) method.

To assess the accuracy of the above semi-automated method for determination of survival curves, the survival curves generated by the lifespan machine were compared to manually-obtained survival curves for four strains of varied virulence and no appreciable difference was observed between lifespans determined automatically compared to manually (Supplemental Figure 8). Virulence assays that involved the use of plasmid-carrying bacterial strains were performed on SK plates supplemented with 20 µM gentamicin.

### Generation of mutant and transgenic *P. aeruginosa* strains

#### Generation of PA14 strains

A PA14*Δcas* in-frame deletion mutant was constructed using a method described previously (Djonovic 2013) that employed a sequence that contained regions immediately flanking the coding sequence of the *cas* genes. This fragment was generated by a standard 3-step PCR protocol using Phusion DNA polymerase (New England Biolabs) and then cloned into the *Xba*I and *Hind*III sites of pEX18A (Prentki and Krisch, 1984), resulting in plasmid pEX18-*CIF*. pEX18-*CIF* was used to introduce the deleted region into the wild-type PA14 genome by homologous recombination. *Escherichia coli* strain SM10 pir was used for triparental mating. The deletion of the Cas genes was confirmed by PCR. For the expression of Cas genes in PAO1, the *P. aeruginosa* PA14 *cas* genes were cloned into the *Hind*III and *Xba*I sites of pUCP19 (West et al., 1994), creating plasmids pUCP-*cas* (referred to as p(Cas+)). The resulting plasmid was transformed into *P. aeruginosa* PAO1 by electroporation to generate the strain PAO1 p(Cas^+^).

#### Generation of z8 strains

Gene deletions in the z8 strain were obtained using the endogenous type I-F CRISPR-Cas present in this strain. In brief, the gentamicin selectable plasmid pAB01 was modified to introduce a spacer targeting the gene of interest and also a homologous recombination (HR) template with arms flanking the genomic region to be deleted (600-800 bp homology arms). The corresponding plasmid so obtained is referred to as ‘editing plasmid’. The z8 bacterial cells were washed twice with 300 mM sucrose and subjected to electroporation (800 ng of editing plasmid, 2 mm gap width cuvettes, 200 Ω, 25 µF, 2500 V using a Gene Pulser XCell machine (Bio-Rad)). All steps were performed at room temperature. Transformants were selected on LB plates with gentamicin 50 µg/mL. Transformant colonies were re-streaked in LB Gentamicin plates and genotyped by PCR. After obtaining the desired genomic modification, the editing plasmid was cured by passage of the strain in liquid LB culture without antibiotic. Plasmid pHERD30T (gentamicin selectable) was used for the expression of genes associated with virulence, gene(s) of interest (with surrounding regulatory sequences) were cloned using Gibson assembly.

### Bacterial growth rates

A random subset of 33 strains that span the virulence range, was used to determine bacterial growth rates. Overnight cultures of each strain (20 µl, O.D. = 1.5-2) were inoculated into 180 µl of LB medium in 96 well plates. The optical densities at 650 nm were measured using the SpectraMax 340 microplate reader (Molecular Devices, CA, USA) every 15 minutes for 33 hours. The experiment was performed at 25°C, the same temperature used for the worm assays, and the plates were shaken for 5 seconds before the measurements by the plate reader to allow aeration. The Softmax Pro 6.2.1 (Molecular devices, CA, USA) software was used to analyze the data. Specific growth rates (µ) were calculated based on the exponential phase of the growth curves. The µ values were calculated using the following formula: OD = N e^*µt*^ where OD is the measured optical density, N the initial optical density, and *t* the time.

### Genomic analysis of *P. aeruginosa* strains

A full list of *P. aeruginosa* species, consisting of 1734 strains, was downloaded from RefSeq database (Tatusova et al., 2016) (on December 2016). In addition, the corresponding annotation files that include (1) genomic sequences, (2) nucleotide and (3) protein sequences for coding genes, and (4) feature tables were downloaded from the RefSeq database as well. Next, several filtration steps were applied to remove strains that: (1) had no proper 16S rRNA annotations (missing sequence, or sequence that is shorter than 1000 nts, or sequence that showed less than 80% identity to PA14 16S rRNA); (2) contained more than 100 core genes with multiple members or were missing more than 15% of the core genes. The second filter was applied after one round of clustering with CD-HIT (Fu et al., 2012) and identification of core genes (see details below). This process resulted in a final set of 1488 strains (Supplemental Table 7).

### Clustering analysis of *P. aeruginosa* coding sequences

The protein sequences of 1488 strains (obtained from the RefSeq database ftp://ftp.ncbi.nlm.nih.gov/genomes/all/GCF/) were clustered using CD-HIT (v4.6.5), with the following settings -c 0.70 -n 5 -g 1 -p 1. The procedure yielded 23,793 clusters of homologous genes. The output of the clustering analysis was post-processed to generate a statistical report that lists for each cluster (*i.e.* each homologous gene) the representative sequence, its function, the total number of occurrences of the gene across the full set of 1,488 strains, and the number of strains that contain at least one copy of the gene. A presence/absence matrix for each gene across 1488 strains was generated. In addition to the full matrix, a presence/absence matrix for the collection of 52 experimentally studied strains was extracted. Gene clusters that had no representatives in these 52 strains were removed, resulting in a matrix with 11,731 genes (Supplemental Table 8).

### Phylogenetic analysis

Core-genes across the 1488 strains were defined as genes present in more than 90% of the strains in a single copy only (resulted in 3494 core-genes). For each cluster representing a core gene the following steps were applied: the corresponding DNA sequences were aligned using MAFFT default parameters (version 7.273) (Katoh and Standley, 2013); gblocks (ver 0.91b) (Castresana, 2000) was applied on the alignment to remove poorly aligned positions (with parameters -t=d -b5=a); an in-house code was used to remove all the invariant positions (excluding gaps); the alignments were padded with gaps for strains in which the core gene was missing. All the alignments were then concatenated to a final alignment of 523,361 nucleotides. The program FastTree (Price et al., 2010), version 2.1, with settings: -gtr, was then used to generate the phylogenetic tree of the 1,488 strains. The interactive Tree of Life web-based tool (Letunic and Bork, 2016) was used for visualization of the resulting phylogenetic tree. Information about MLST, source (clinical/environmental) and strains that are part of the experimental collection was incorporated into the tree view. A phylogenetic tree of the 52 experimentally studied strains was extracted from the phylogentic tree of the 1488 strains using the ‘ape’ package in R.

### Statistical test for association of genetic elements (coding/non-coding genes) with virulence

The Mann-Whitney (MW) ranking test and linear-regression (LR) analysis were applied to every gene to test the association of the presence/absence pattern with virulence. Genes were considered associated if both tests yielded a p-value lower than 0.05, and at least one of the tests yielded a p-value smaller than 0.01. Among the virulence-associated genes, genes with negative slope (based on linear regression) were associated with low survival/high virulence (referred to as high-virulence associated or HVA), while genes with positive slope were associated with high survival/low virulence (referred to as low virulence associated or LVA). All the p-values are shown in log10 scale as absolute values. The reliability of the p-values was assessed using a permutation test as described below.

### Permutation test to assess the reliability of the p-values

10,000 permutations of the virulence values and their assignment to strains were generated (*i.e*. median worm survival values) and the MW and LR association tests were repeated for each permutation. Then, for each gene the number of times that it received a better p-value using the shuffled virulence data compared to the original one was recorded, separately for MW and LR. The reliability score was calculated by dividing the above count by 10,000. The MW and LR p-values were considered reliable if their reliability score was less than 0.05.

### Assessment of confounding effects due to population structure

The phylogenetic method reported by Collins and Didelot (2018), known as treeWas, was used to address the potential influence of population structure in the statistical association between accessory genes and virulence. The method was applied on the input consisting of: (1) 11,731 gene clusters presence/absence matrix, (2) median survival vector and (3) phylogenetic tree of the 52 strains. The method returns as output three types of scores and their corresponding p-values for every gene cluster: (1) “Terminal Score” which measures sample-wide association between genotype (gene presence) and phenotype (median survival), without relying on the phylogenetic tree; (2) “Simultaneous Score” which measures the degree of simultaneous change in the phenotype and genotype across branches of the phylogeny; and (3) “Subsequent Score” which measures the proportion of the tree in which genotype and phenotype co-exist. The computed scores were considered significant if their p-values < 0.05. (Supplemental Table 2).

### Collection of known non-coding RNA (ncRNA) in *P. aeruginosa*

The collection of ncRNAs (excluding rRNAs and tRNAs) in *P. aeruginosa* was constructed using two resources: RFAM 12.2 (Nawrocki et al., 2015) and RefSeq annotations (Tatusova et al., 2016). First, 75 non-coding RNA families were extracted from RFAM, with a total of 1,363 sequences across *P. aeruginosa* strains. To get the representative sequences (there could be more than one) for each family, the sequences of each family were clustered using CD-HIT-est (with 80% identity). This analysis resulted in 115 sequences (representing 75 different ncRNA families). Second, using RefSeq annotations of the 1,488 strains, 2,549 ncRNA sequences were extracted. Blasting these sequences against RFAM families followed by clustering analysis revealed additional 8 families that are missing from RFAM. All together our collection comprised of 83 ncRNA families, represented by 123 sequences. Finally, the collection of the 123 sequences was blasted against the 1,488 genomic sequences, and a presence/absence matrix for each of the sequences in all the strains was generated. Rows that represent sequence members from the same family were collapsed, resulting in matrix with 83 rows.

### Collection of previously identified virulence genes in *P. aeruginosa*

A list of virulence genes, in either PA14 or PAO1, was downloaded from (Bartell et al., 2017). The list was filtered to contain only genes that were reported to contribute to *P. aeruginosa* virulence towards *C. elegans*, resulting in 56 genes. Another four genes were added based on the publication (van Tilburg Bernardes et al., 2017). The homologous gene clusters that contained the above genes were marked as virulence genes. The full list of 60 virulence genes is found in Supplemental Table 3.

### Analysis of CRISPR-Cas systems

#### Identification of CRISPR-Cas systems

The presence of CRISPR-Cas systems in the genomes of our *P. aeruginosa* collection was determined by identifying the gene clusters that encode for Cas proteins.

#### Identification of anti-CRISPR genes

The most up to date collection of anti-CRISPR genes was downloaded from (Marino et al., 2018), consisting of 41 sequences (https://tinyurl.com/anti-CRISPR). Annotations (*e.g*. CRISPR-Cas subtype inhibited) for each sequence were maintained. The representative sequences of the clusters of homologous genes (see CD-HIT clustering above) were blasted against the anti-CRISPR sequences using blastp (Altschul et al., 1997) and e-value threshold of e-10. A coverage of more than 35% of the anti-CRISPR sequence, was considered a hit.

#### Defining active/inactive systems

The annotation on the type of CRISPR-Cas system(s) that is inhibited by each anti-CRISPR protein was used to define CRISPR-Cas activity. The type(s) of CRISPR-Cas systems of every strain were matched to the type(s) inhibited by the anti-CRISPR genes present in the same genome. Strains where all present CRISPR-Cas system(s) are inhibited by type-matching anti-CRISPR proteins were considered inactive.

#### CRISPR Spacer arrays collection

The collection of CRISPR spacer sequences across all 1,488 strains was generated by applying the CRISPR Recognition Tool (CRT1.2-CLI.jar) (Bland et al., 2007) on genomic sequences, with default parameters. Since the tool works only with single fasta records, the genomic sequences (contigs and scaffolds) of each strain were merged before the application of the tool, and then the results were mapped back to the original sequences using an in-house code. A total of 35,340 spacer sequences were identified (some sequences were present more than once in the collection).

#### Targets of CRISPR spacers on P. aeruginosa pangenome

The program blastn (Altschul et al., 1997), with default parameters was used to identify matches for the full spacer’s collection against the DNA sequences of all protein coding genes. The homologous gene clusters that contained the targeted genes were marked as CRISPR targets. The above set of targets and spacers was further filtered, and spacers where its target is located in the same genome were tagged as ‘self-targeting’ spacers. In order to use self-targeting spacers to estimate CRISPR-Cas ‘inactivity’ an additional criterion was included: the target (protospacer) should be conductive to CRISPR-Cas cutting of the bacterial DNA, i.e. a full spacer-target alignment with PAM presence should exist. A strain was considered CRISPR-Cas ‘inactive’ by the presence of a CRISPR-Cas locus and at least one spacer satisfying the above criterion.

### Analysis of restriction modification (RM) systems

Sequences of RM systems and their type classification were downloaded from REBASE (The Restriction Enzyme Database) (Roberts et al., 2015). The representative sequences of the clusters of homologous genes (see CD-HIT clustering above) were blasted against the RM sequences using blastp and e-value threshold of e^−10^. Several filtration steps were ten applied before marking a gene cluster as an RM gene. Gene clusters were excluded if: (1) the coverage of the RM sequence by the representative sequence was less than 35%; (2) if the gene cluster represents a core gene and (3) the function associated with the gene cluster is not diagnostic to an RM system (*e.g.* permease, topoisomerase). 227 gene clusters passed the criteria.

Next, the RM genes of every strain, were extracted and re-ordered based on their genomic location. Using the location of the genes, ‘gene blocks’ were determined as groups of genes separated by less than 8 intervening genes.

For every gene, the best matching RM component from REBASE was used to assign an RM type (either type I, II, III or IV) and identity the RM component (methylase, nuclease, specificity factor, etc). Every gene with a match to a type IV RM was established as a type IV system.

Next, all other RM systems (types I to III), were defined based on the presence of methylase genes. A gene singleton (*i.e*. not belonging to any gene block) matching a type II methylase, was established as type II RM system. RM systems inside gene blocks were assigned based on the following criteria: (**a**) 1 or 2 methylases must be present per RM system; (**b**) all gene components of a given RM system, congruently match a single type of RM system. To assess the quality of our RM data, we compared our predictions to REBASE data. Seven strains from our collection have their genomes annotated in the REBASE website. 4 strains have the exact same number of RM systems, while the RM count of the 3 remaining strains differ by one RM. No statistical difference exists between our method and REBASE with regard to the RM count of strains (chi square test, p = 0.18).

### Analysis of novel defense systems

Protein accession numbers belonging to ten novel defense systems were downloaded from (Doron et al., 2018) and were filtered to keep only *P. aeruginosa* proteins. Each protein sequence was annotated with system type and specific system component. The protein sequences were then extracted from RefSeq. The representative sequences of the clusters of homologous genes (see CD-HIT clustering above) were blasted against the protein sequences using blastp (Altschul et al., 1997) and an e-value threshold of e^−10^. A filtration step was applied before marking a gene cluster as a defense system gene. Gene clusters were excluded if: (1) the coverage of the defense system sequence by the representative sequence was less than 35%. Next, the candidate genes for novel defense systems of every strain, were extracted and re-ordered based on their genomic location. Using the location of the genes, ‘gene blocks’ were determined as groups of genes separated by less than 8 intervening genes. All novel defense systems, were defined based on the presence of a set of 2 or more genes uniformly matching a variant of the novel systems as reported by (Doron et al., 2018).

---

**Supplemental Figure 1.**
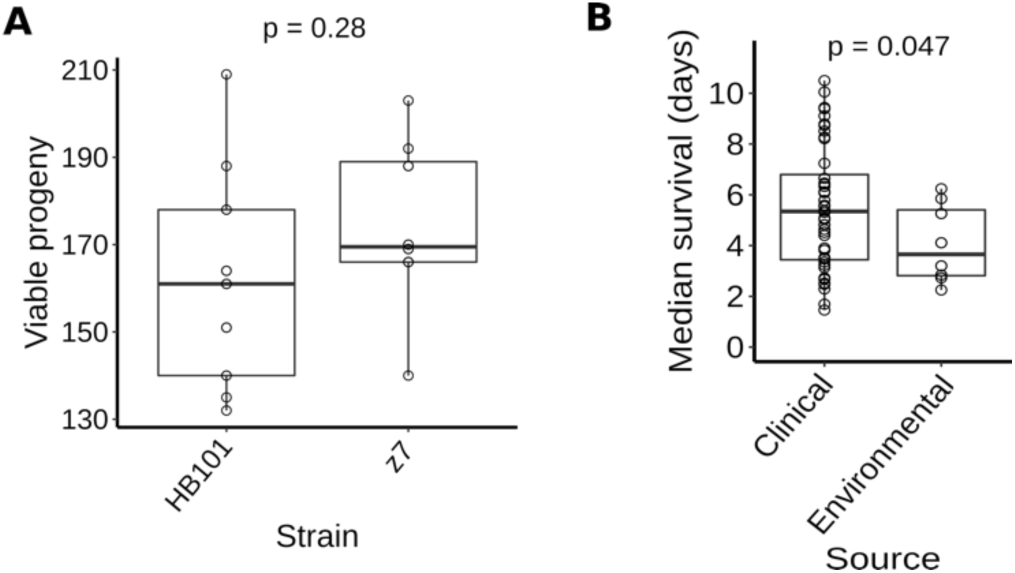
Traits of the interaction between *C. elegans* and *P. aeruginosa* isolates. **A**) Viable progeny counts for *P. aeruginosa* z7 and *E. coli* HB101. Adult *C. elegans* hermaphrodites were exposed to the above-mentioned bacterial strains using the same conditions for virulence assay with the exception that no FUDR was added (SK plates, 25°C). The total progeny of individual worms was manually counted. Comparison of the two conditions was done using the Welch *t*-test (p-value indicated). **B)** Box plot of worm median survival in relationship with strain source (environmental or clinical). p-value is indicated for the Welch *t*-test comparison of virulence (*i.e.* induced worm median survival) between clinical and environmental strains.

**Supplemental Figure 2.**
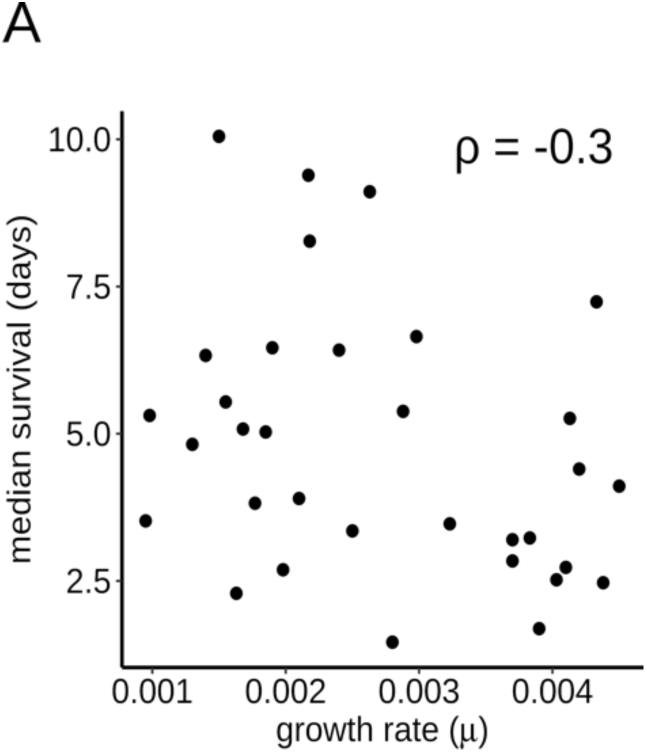
Relationship between bacterial growth rate and virulence. Association between bacterial growth rates in LB medium (µ) and virulence among *P. aeruginosa* strains (median survival in days).

**Supplemental Figure 3.**
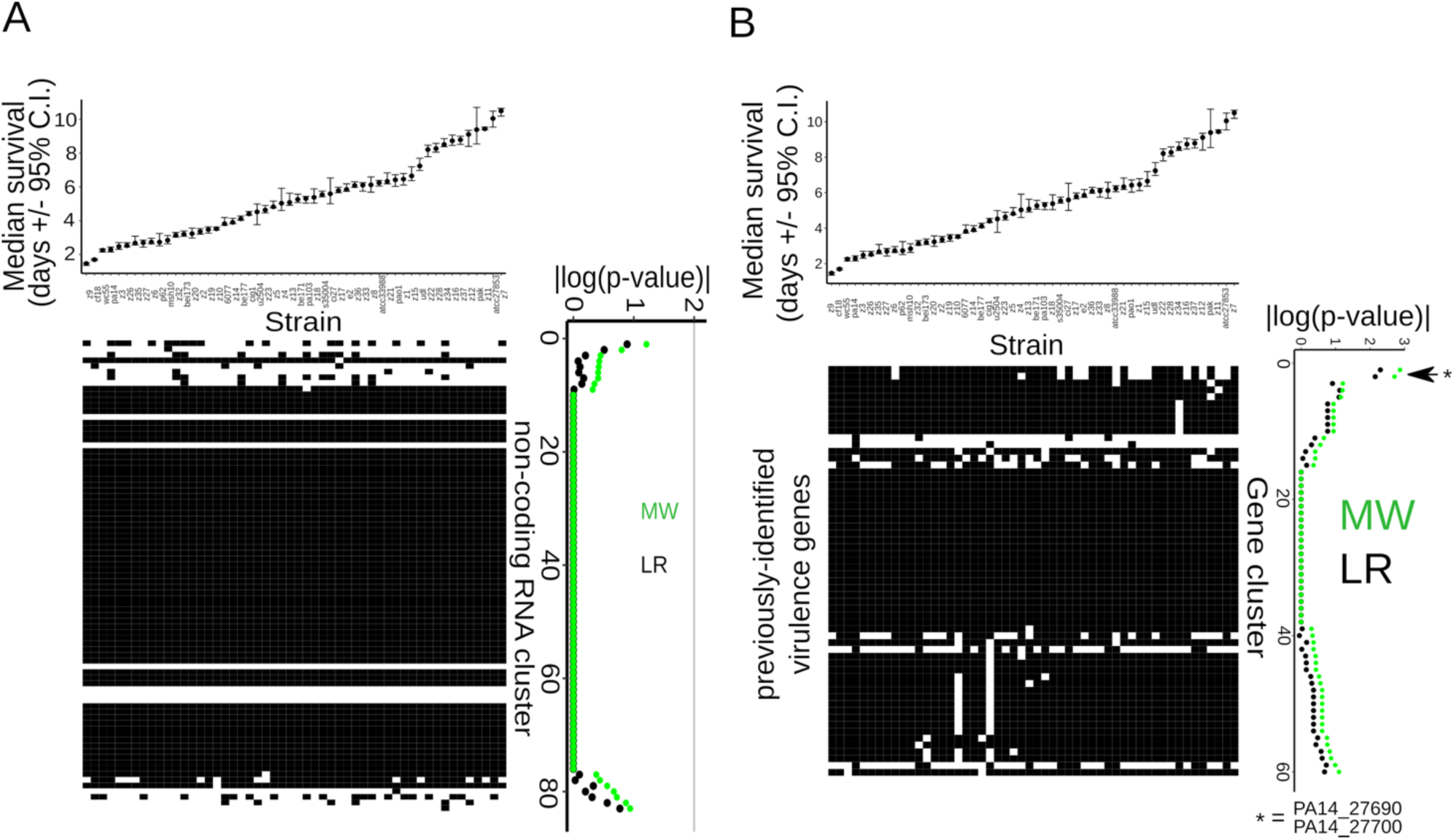
Association between non-coding RNA and previously known virulence genes and virulence. **A**) Association between non-coding RNAs of *P. aeruginosa* and bacterial virulence: (Top panel) Median survival of adult *C. elegans* worms, similar to Figure 2B. (Bottom left panel): gene presence/absence matrix for non-coding RNAs. Presence is indicated with black squares and absence with white squares. Non-coding RNAs (rows) are aligned with the corresponding MW and LR p-values (bottom right panel), shown as | log_10_(pval)|. Rows are ordered from association with high virulence to association with low virulence. **B**) Distribution and association of previously known virulence genes. (Top panel) Median survival of adult *C. elegans* worms exposed to the studied collection of *P. aeruginosa* strains. The strains are ordered from high to low virulence (left to right) and aligned with the matrix below. (Bottom left) gene presence/absence matrix for known virulence genes. Gene presence is indicated with black squares and absence with white squares. Genes (rows) are aligned with the corresponding p-values. (Bottom right) Association statistics (p-value of MW and LR tests) for the genes (shown as | log10(p-value)|). Rows are ordered from association with high virulence to association with low virulence.

**Supplemental Figure 4.**
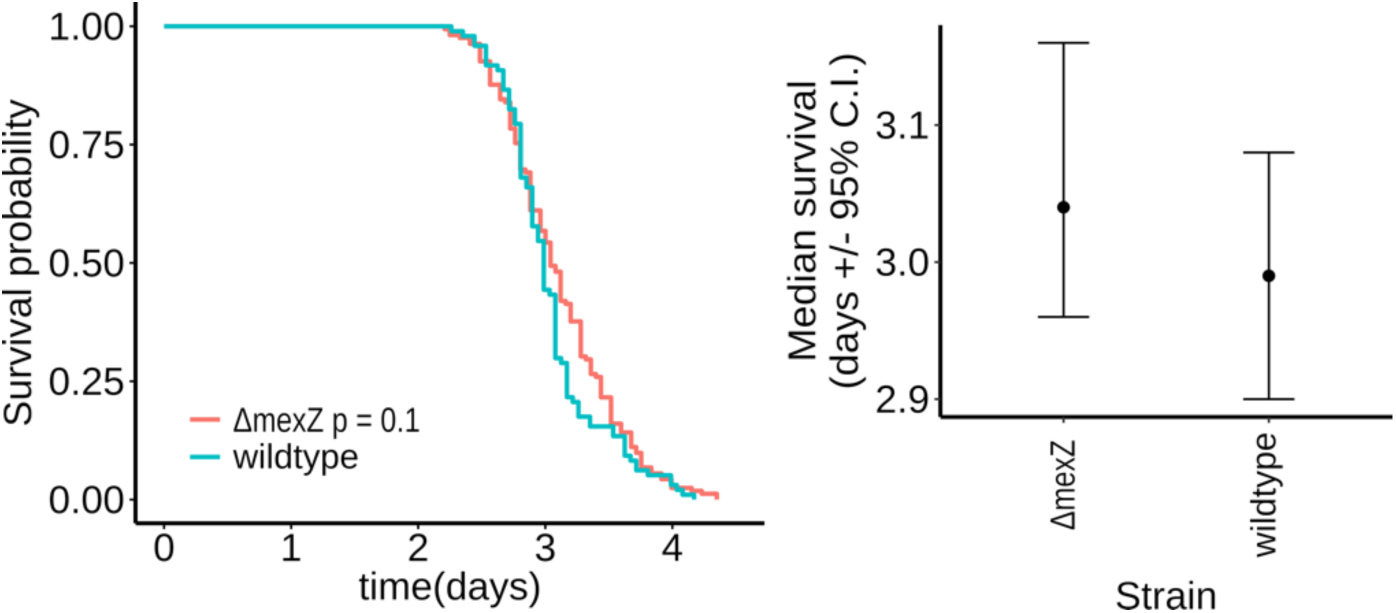
Bacterial virulence upon loss of mexZ gene. Survival curves (left panel) and median survival (right panel, confidence interval ‘C.I.’) of adult *pmk-1(lf) C. elegans* worms exposed to wild-type and *ΔmexZ* strains of *P. aeruginosa* z8. Pairwise comparison of the survival curves between the two strains was done using the logrank test. The test p-value is indicated in the curve legend.

**Supplemental Figure 5.**
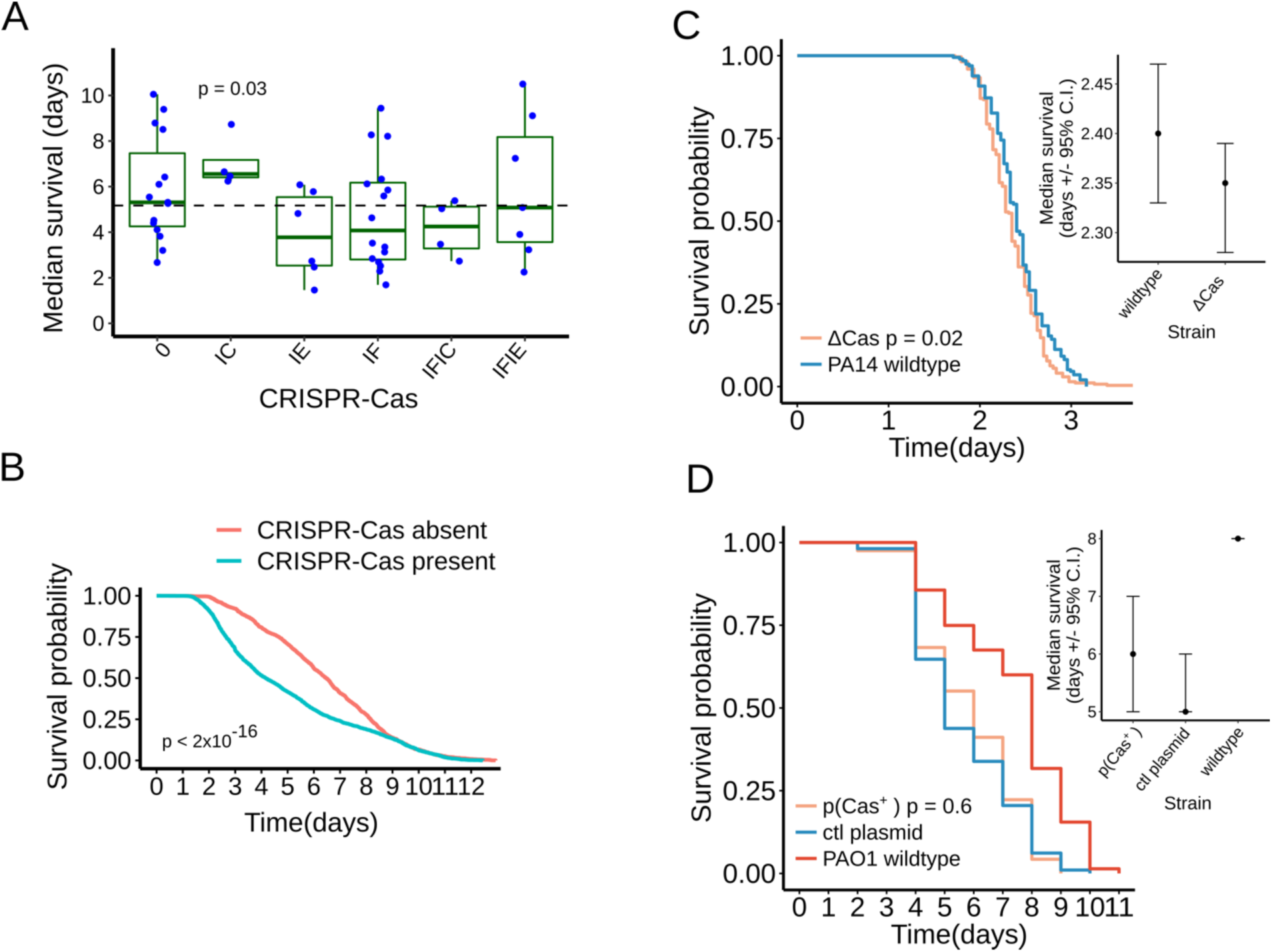
Relationship between CRISPR-Cas systems and virulence. (**A**) Relationship between CRISPR-Cas subtypes and virulence. Strains are categorized by their combination of CRISPR-Cas subtypes. Strains with type CRISPR-Cas I-C systems have significantly lower virulence than their complementary strain set (Welch t-test, p-value = 0.03). (**B**) K-M Survival curves of adult *C. elegans* worms exposed to the studied collection of 52 *P. aeruginosa* strains partitioned according to presence (in cyan color) or absence (in red color) of host CRISPR-Cas systems. Survival data for the pooled strains were aggregated as described in Materials and Methods. The p-value of a long-rank test between the two subgroups is indicated. (**C-D**) Survival curves (left panels) and median survival (right panels, confidence interval ‘C.I.’) of adult *C. elegans* worms exposed to strains of *P. aeruginosa*. (**C**) Virulence of PA14 wildtype and PA14 with deletion of the type I-F Cas genes (ΔCas). (**D**) Virulence of PAO1 wildtype; PAO1 with plasmid expressing the type I-F Cas genes (pCas^+^); PAO1 with control plasmid (ctl plasmid). Pairwise comparison of the survival curves was done using the logrank test. The p-values are indicated in the respective legend.

**Supplemental Figure 6.**
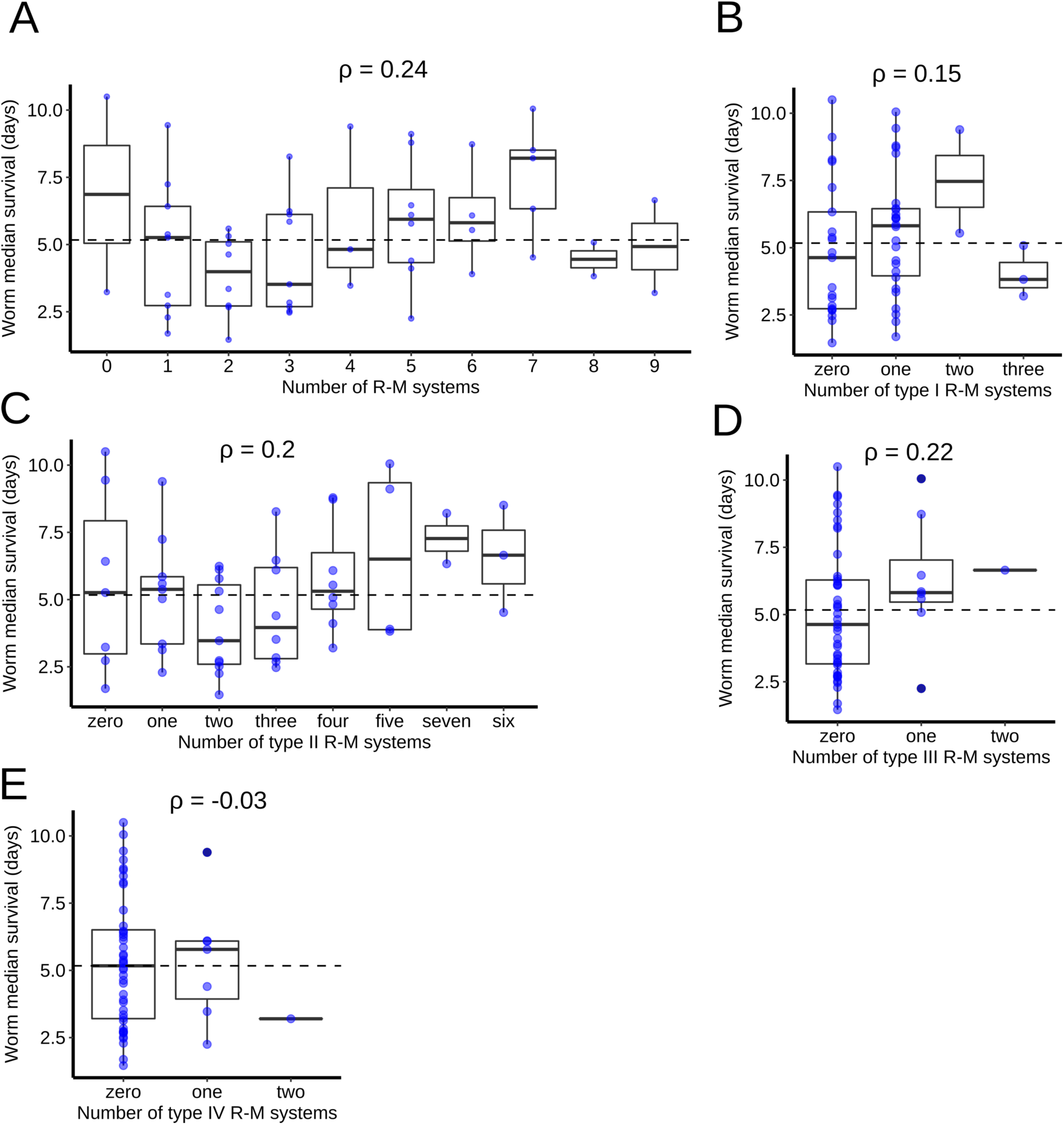
Relationship between Restriction-Modification (RM) systems and virulence. **A-E)** Box plots of worm median survival (virulence) in relationship with the abundance and type of RM systems. **A**) The total number of RM systems per strain is displayed. **B-E**) The number of RM systems per strain is displayed separately for type I (**B**), II (**C**), III (**D**) and IV (**E**) systems. Correlation values are indicated in all graphs (r, Spearman rank correlation). The median virulence of the complete set of strains displayed on each graph is indicated with the dashed horizontal line.

**Supplemental Figure 7.**
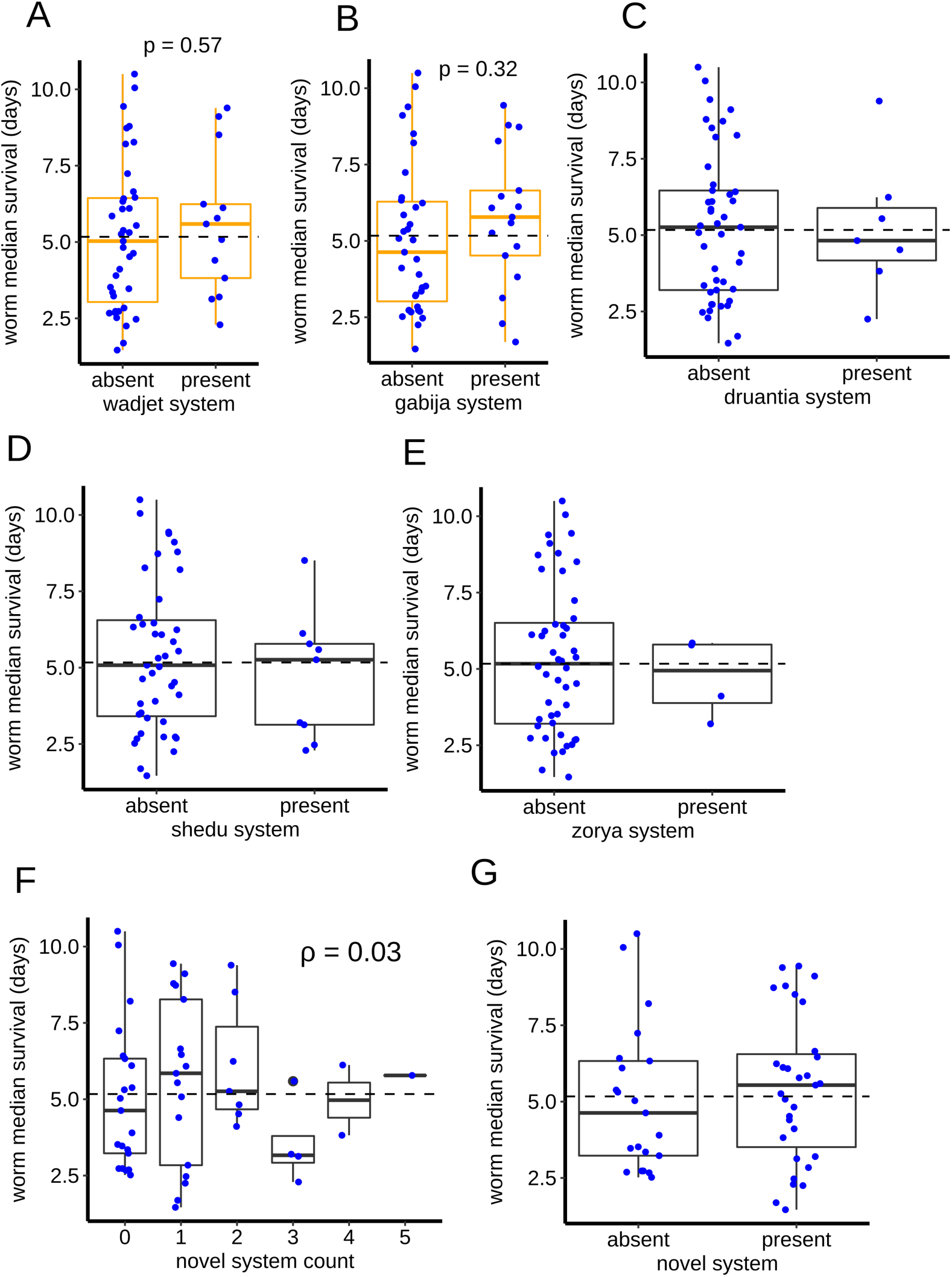
Relationship between recently described defense systems and virulence. (**A-E**) Box plots of worm median survival in relationship with novel defense system abundance and types. The presence/absence of six novel systems in relationship with median worm survival, displayed separately for wadjet (**A**), gabija (**B**), druantia (**C**), shedu (**D**) and zorya (**E**) systems. **F)** The total number of novel systems per strain is displayed. **G**) The presence/absence of novel defense systems is displayed. In all graphs, no difference in virulence compared to their complementary strain sets is observed (Welch t-test, all p-values > 0.05). The median virulence of the complete set of strains displayed on each graph is indicated with the dashed horizontal line.

**Supplemental Figure 8.**
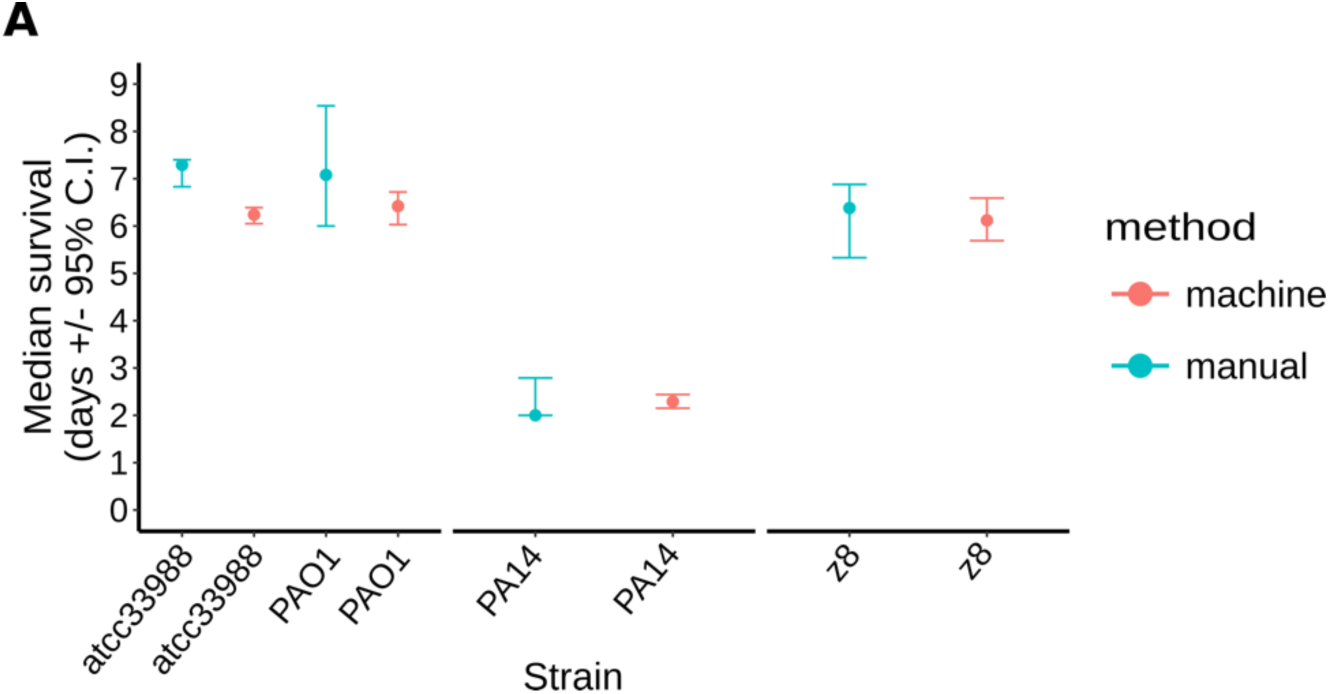
Comparison between two methods to determine median survival. Median survival of adult *C. elegans* worms exposed to four *P. aeruginosa* strains (confidence interval, C.I.) scored with two distinct methods. The methods to obtain the median survival estimates are: semi-automated scanning procedure (referred to as ‘machine’); manual scoring with a pick (referred to as ‘manual’).

## ACKNOWLEDGEMENTS

We would like to acknowledge members of the Ambros and Mello laboratories for feedback about this research project. We would also like to thank Deborah McEwan for assistance with the lifespan machine, Zeynep Mirza for contributing the growth rate measurements, Joseph Bondy-Denomy for sharing plasmid reagents and Veronica Kos for help with strain requests. Some of the investigated bacterial strains were obtained from International Health Management Inc. Regarding *C. elegans*, some strains were provided by the CGC, which is funded by NIH Office of Research Infrastructure Programs (P40 OD010440).

## AUTHOR CONTRIBUTIONS

A.V.R, I.V-L, V.A designed the study, performed and analyzed experiments, wrote the manuscript. Z.C. constructed *P. aeruginosa* strains, Z.C. F.A. reviewed the results and the manuscript.

## SUPPLEMENTARY TABLE LEGENDS

**Supplemental Table 1**. Description and features of the experimentally studied collection of 52 *P. aeruginosa* strains. The 52 strains experimentally studied strains are listed, altogether with all the features derived from this study.

**Supplemental Table 2**. Genes significantly associated with virulence. Description of the 79 genes that comprise the HVA and LVA sets.

**Supplemental Table 3.** Known virulence genes in the interactions between *P. aeruginosa* and *C. elegans* under SK condition

**Supplemental Table 4. Nomenclature for the experimentally studied bacterial genes**

A set of genes associated with virulence are termed for the *P. aeruginosa* strains z8 and PAO1. Genes that constitute a gene block frequently found in multiple tandem copies in various strains are termed teg(G to N), for ‘tandem element gene’. The region encompassing from tegG to tegN is referred to as ‘teg gene block’. The Refseq gene ‘NT41_RS12090’ is termed ghlO (glycosyl hydrolase like ORF) as it exhibits similarity to domain Cdd:cd06549 (E-value: 0.02, CDD database). The PAO1 genes: PA2228, vqsM, qsrO, and PA225, constitute a putative operon (Köhler et al., 2014) that is referred to as ‘qsr’ operon.

**Supplemental Table 5. Bacterial strains generated in the present study**

Strains generated in the present study are described with a strain name (AVPae #) and genotype (in both full and short formats).

**Supplemental Table 6.** Gene targeted by CRISPR spacers.

**Supplemental Table 7**. Description of *in silico* studied set of 1448 *P. aeruginosa* strains

**Supplemental Table 8**. Gene clustering analysis for the *in silico* studied *P. aeruginosa* strains. Shown are only gene clusters that contain sequences from the studied 52 strains.

